# Molecular trick to reverse S_N_2 mechanism in hydrolytic enzyme

**DOI:** 10.1101/2025.09.17.676745

**Authors:** Martin Toul, Sergio M. Marques, Tadeja Gao, Hana Bernhardova, Ondrej Vavra, Veronika Novakova, Jiri Damborsky, David Bednar, Zbynek Prokop, Martin Marek

## Abstract

Hydrolytic haloalkane dehalogenase enzymes catalyze an S_N_2 nucleophilic substitution to erase halogen substituents in organohalogen compounds. The acid-base-nucleophile triad secures irreversible S_N_2 displacement of the halogen for the hydroxyl derived from the water. Catalysis relies on the protonatable imidazole ring of the histidine base, and its substitution with an asparagine traps the enzyme in a covalently bound intermediate state, a principle exploited in the widely used HaloTag technology. In contrast, the histidine-to-phenylalanine substitution triggers reversibility of the S_N_2 mechanism, but the molecular trick by which it reprograms the catalytic pathway remains unknown. Here, we show that the phenylalanine at the site of the histidine base spatially disturbs the adjacent residues, leading to the remodeling of surrounding active-site loops. Consequently, rerouting of the access tunnels imparts distinctive kinetic behavior, featuring a reversible S_N_2 chemical step that facilitates transhalogenation reactions. This information is crucial for engineering next-generation biocatalysts for sustainable chemistry.

**Highlights:**

- Catalytic triad (glutamate-histidine-aspartate) secures irreversible S_N_2 mechanism
- Histidine-to-phenylalanine substitution in the triad leads to a reversible S_N_2 reaction
- Enzyme active site rerouting is a hallmark of the catalytic reprogramming
- Basis for designing biocatalysts for sustainable transhalogenation chemistry

**The bigger picture:** Haloalkane dehalogenases catalyze the hydrolytic cleavage of the carbon-halogen bond in halogenated hydrocarbons, a feature used in a wide variety of industrial and biotechnological processes. In HaloTag technology, the catalytic histidine is replaced by an asparagine to enable a stable covalent bonding between a probe and a protein for biological imaging, affinity purification, etc. Remarkably, this originally irreversible S_N_2 process becomes fully reversible in a histidine-to-phenylalanine mutant. Moreover, halogen ion product is released at this stage, offering a greener alternative for transhalogenation reactions compared to the conventional synthetic approach and a way to recycle environmentally harmful halogenated compounds. However, the optimization of enzymes is needed for applications on an industrial scale. Therefore, we investigated the kinetics of the S_N_2 step and structural features of these two enzyme mutants, providing a basis for subsequent optimization efforts.

## Introduction

Enzymes are highly efficient catalysts needed for numerous reactions occurring in living organisms, biotechnology, and the pharmaceutical industry.^1^ Because of their catalytic specificity and environmentally friendly nature, they are employed in synthetic chemistry, enabling the production of complex molecules with high selectivity and a greener approach compared to conventional synthetic methods.^2–3^ Nevertheless, their catalytic properties must be tailored to meet the needs of industry and research. Often, site-directed mutagenesis suffices for these needs, with mutations usually done in or close to the catalytic site.^4–6^ The identity of an amino acid that will replace the original one should be carefully chosen,^7–8^ since it can cause a long-distance effect outside of the active site.^9–11^ When structural information about the target enzyme is limited or unknown, directed evolution can be a valuable tool in protein engineering, though time and energy-consuming. Ideally, when a detailed understanding of the structural basis related to enzyme function is known, rational design can save time and energy in the discovery of novel biocatalysts.^12^

Case in point is a family of haloalkane dehalogenases (HLDs; EC 3.8.1.5), microbial enzymes used in many industrial and biotechnological processes.^13^ They employ the mechanism of S_N_2 nucleophilic substitution to catalyze the hydrolytic cleavage of the carbon-halogen bond in halogenated hydrocarbons to the corresponding alcohol, halide ion, and a proton (**Figure 1A**).^14^ The conserved catalytic pentad consists of five residues: an aspartate nucleophile, a histidine base, aspartic or glutamic acid, and two halide-stabilizing residues, mostly a tryptophan-tryptophan or tryptophan-asparagine pair.^14–15^

**Figure 1:**
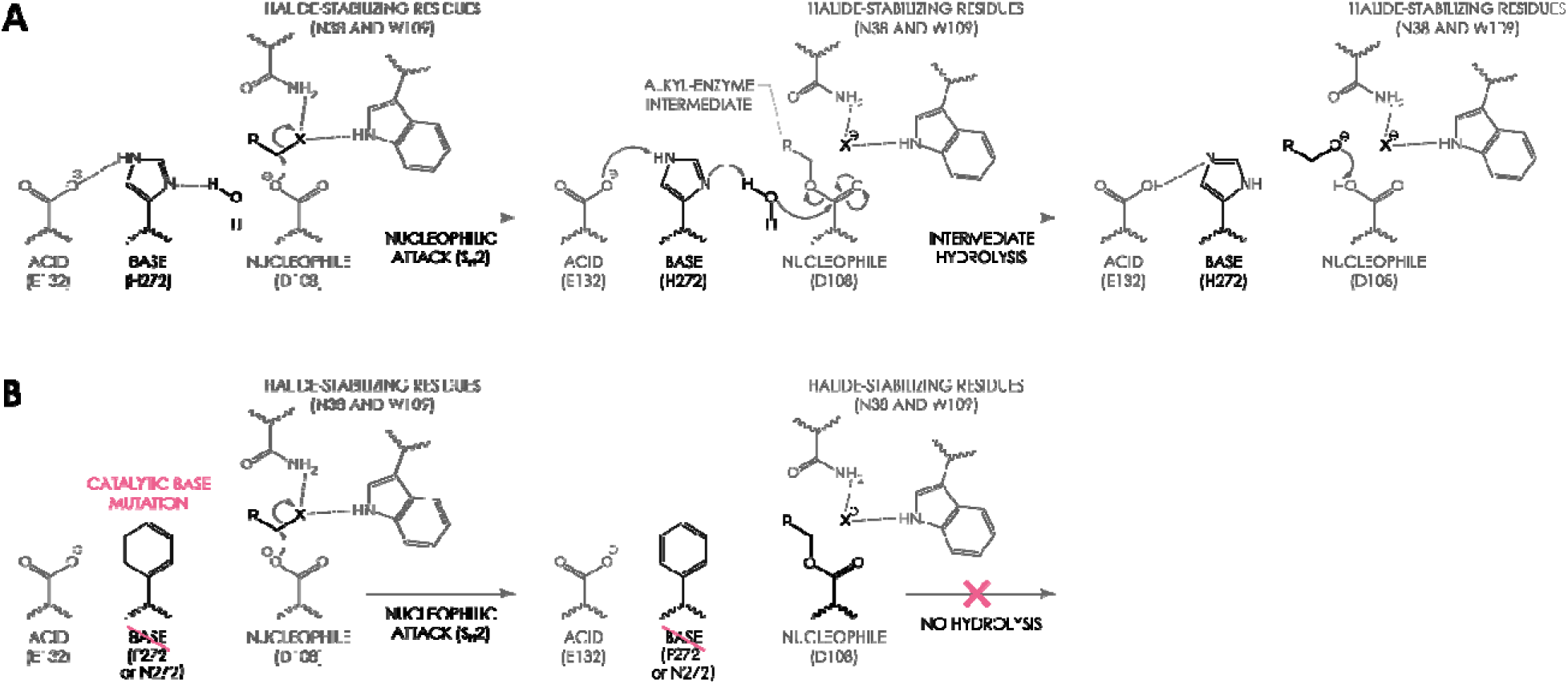
Schematic representation of the hydrolysis step of the dehalogenase reaction catalyzed by HLDs. (A) Wild-type HLDs contain catalytic histidine (H272 in the case of LinB), which catalyzes the hydrolysis of the alkyl-enzyme intermediate. (B) The mutation of the catalytic histidine to phenylalanine or asparagine (H272F/N in the case of LinB) blocks the hydrolysis step.

Substitution of the catalytic histidine enables the usage of HLDs in protein tagging (so-called HaloTag technology). First HaloTag versions used the histidine-to-phenylalanine mutation.^15^ The later experiments showed that the histidine-to-asparagine substitution is a better choice in terms of stability;^16–17^ confirmed in numerous crystallographic studies.^18–22^ Here, the protein of interest is genetically fused with an HLD, which is covalently bound to a synthetic ligand, bearing the desired property (e.g., fluorescence). To enable this covalent bond, the hydrolysis of the HLD catalytic cascade needs to be disrupted, which is achieved by a mutation of the catalytic histidine to asparagine (**Figure 1B**).^23–24^ HaloTag nowadays covers various applications, including protein immobilization,^25^ purification,^26^ cellular protein imaging,^27–28^ imaging *in vivo*,^29^ etc.

On the other hand, the histidine to phenylalanine mutation of the same HLD causes an opposite effect. The stable covalent modification that is formed in the asparagine mutant becomes substantially weaker in the phenylalanine mutant. This enables the reversibility of the S_N_2 step, a desired property for transhalogenation reactions, contributing to a greener approach in synthetic chemistry. Beier *et al*.^24^ reported the transhalogenation potential of several HLD members with histidine to phenylalanine mutation. However, optimization of these mutants is needed to target industrial applications. Efficient optimization can be achieved by rational design which requires structural information on HLD mutants.

To this end, we studied the effect of a single-point mutation of the catalytic histidine to phenylalanine or asparagine on the overall protein function and structure by transient kinetic experiments, X-ray crystallography, and molecular dynamic simulations. As a model, we used LinB from *Sphingobium japonicum* UT26, a member of the HLD family, where three LinB variants (wild type, H272F, and H272N) were studied. We present how differences seen in the activity of all three enzymes can be justified by structural changes seen by X-ray crystallography. We observed an unexpected structural change in the H272F mutant resulting in the opening of a new tunnel, which is important for a small molecule transfer and was never observed in the HLD family before. The combination of X-ray structure and MD simulations provided an explanation for why the tunnel opening occurs and provides the link between structure and catalytic behaviors of these variants.

## Results

### Ligand binding and catalytic potential

We performed transient kinetic experiments for wild-type LinB (LinB-wt), and two mutants LinB-H272F and LinB-H272N at two pH values 7.5 and 10.5. We employed 1-chlorohexane as a substrate and Cl^-^ ion as a product. Kinetics data were collected by monitoring the changes in the native tryptophan fluorescence upon binding and unbinding of Cl^-^ to catalytic W109.

The elementary steps of the reaction between enzyme (E) and the substrate (R-Cl) are schematically depicted in **Figure 2A**. The reaction steps are the following: binding of the substrate → S_N_2 reaction → intermediate hydrolysis → dissociation of the products. Initially, the substrate binds to the protein via interactions between the chlorine atom and the catalytic residues N38 and W109 (E.R-Cl). This step is followed by nucleophilic substitution, S_N_2, where the chlorine is detached, and the substrate forms an ester bond with the carboxylic group of the catalytic aspartate D108 to yield alkyl-enzyme intermediate (E-R.Cl^-^). Meanwhile, the displaced chloride stays bound to N38 and W109 until the very last step. The third step, which is blocked in the case of LinB-H272F and LinB-H272N, is the intermediate hydrolysis involving the imidazole ring of H272. The imidazole ring activates a water molecule by deprotonation, allowing the hydrolysis of the intermediate and regeneration of the free acidic D108. As a result, the ester bond of the alkyl-enzyme intermediate breaks, yielding the ternary complex of the enzyme with both the alcohol and the chloride product temporarily bound in the active site (E.R-OH.Cl^-^). Finally, the products, alcohol, and chloride, are released from the protein core (E + R-OH + Cl^-^).

**Figure 2:**
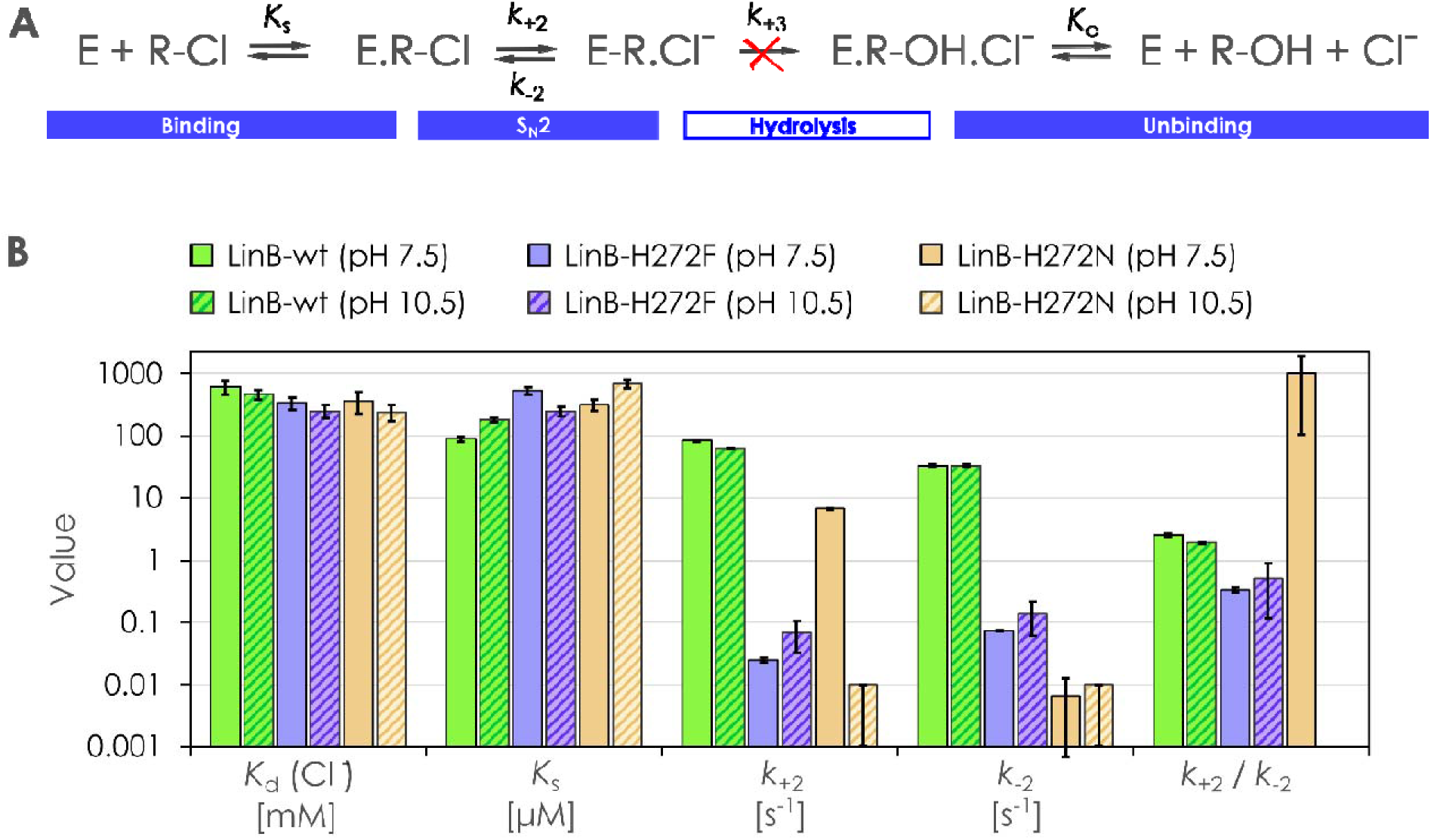
Transient kinetic analysis of the LinB variants. (A) Schematic representation of the binding, two chemical steps (S_N_2 and hydrolysis) and the unbinding step. The red cross symbolizes the inability of the catalyti histidine mutants H272F/N to hydrolyze the intermediate E-R.Cl^-^ and to form the final products. (B) Kinetic constants obtained after mixing Cl^-^ ion or 1-chlorohexane with LinB proteins: LinB-wt (green bars), LinB-H272F (purple bars), and LinB-H272N (wheat bars). Solid bars present results at pH 7.5 and hatched bars at pH 10.5. The values are represented as best-fit values ± standard error values based on nonlinear curve fitting.

The first step, the binding of 1-chlorohexane (substrate) and Cl^-^ ion (product) to the enzyme, shows no substantial differences across LinB variants and pH values. The values of dissociation constants *K*_d_ (for Cl^-^ ion) and *K*_s_ (for 1-chlorohexane) are summarized in **Figure 2B** and **Table S1**. The detailed analyses are depicted in **Figures S1**-**3**. The overall results demonstrate that the Cl^-^ and 1-chlorohexane binding process is affected neither by the mutations nor the pH values.

However, the rates of the second S_N_2 step (conversion of the 1-chlorohexane substrate to the alkyl-enzyme intermediate; E-R.Cl^-^) differ drastically depending on the mutation and pH (**Figure 2B** and **Table S1**). Namely, in LinB-H272F, the rate constants *k*_+2_ and *k*_-2_ are approximately 1,000-fold lower, showing highly impaired kinetics of the E-R.Cl^-^ formation (S_N_2 reaction) compared to LinB-wt. In contrast, LinB-H272N exhibits only a 10-fold lower rate of the chemical step compared to LinB-wt, and more importantly, this chemical step is nearly irreversible (high *k*_+2_/*k*_-2_ ratio), explaining the importance of the specific histidine substitution with asparagine over phenylalanine for a stable HaloTag labeling.

Additionally, the significant influence of pH value on the S_N_2 step is observed for both mutants, while LinB-wt is unaffected. The alkaline pH severely impairs the kinetics of the LinB-H272N, which becomes more similar to LinB-H272F at pH 7.5, probably due to the changes in residue protonation states and modified interactions within the protein. Owing to the prolonged 1-chlorohexane processing by LinB-H272N at pH 10.5 and no detectable exponential fluorescence decrease within the measured time window (100 seconds), only upper limits of the chemical step rate constants *k*_+2_ and *k*_-2_ (0.01 s^-1^) were estimated. In the case of LinB-H272F, the increase in pH does not change the rates of the S_N_2 step, but a new kinetic pattern appears (detailed analysis in **Supplementary Note 1** and **Figure S4**), suggesting the ability of the protein to release the Cl^-^ product at the level of the intermediate. The binding and catalytic pattern of LinB-H272F at pH 10.5, including the reversibility and product release during S_N_2, reveals its potential in the transhalogenation reaction.^24^

### Backbone movements upon a histidine-to-phenylalanine change

To understand the different performance in the activity of LinB variants, we determined the crystal structure of LinB-H272F with a 1.55 Å resolution. The asymmetric unit contains one enzyme molecule that packs well and has good deviations from the ideal geometry and statistics (**Table S2**). The LinB-H272F adopts a canonical HLD fold, with several as-yet-unseen structural features (**Figures 3** and **S5**).

**Figure 3:**
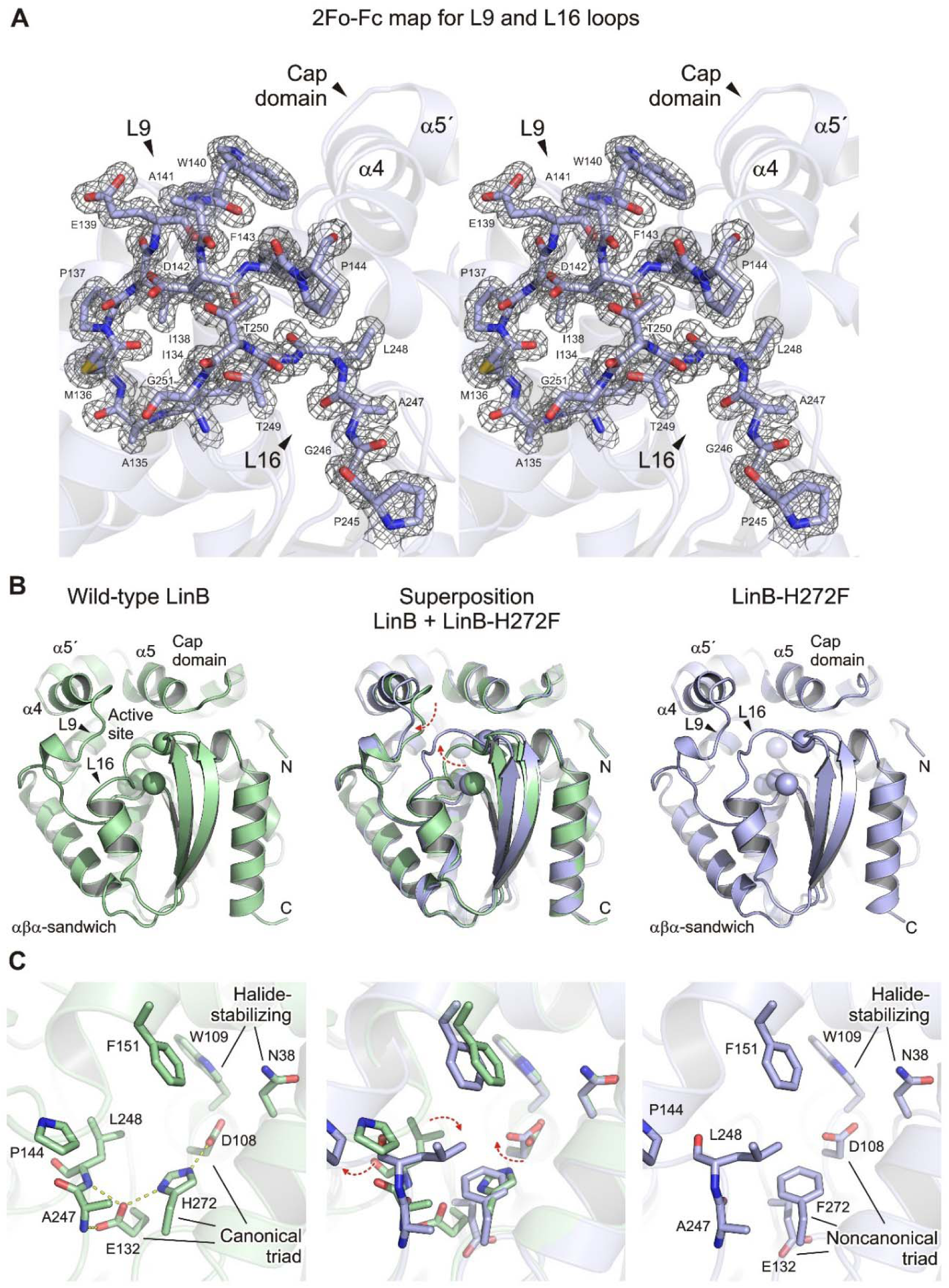
Structural comparison between the wild-type LinB (LinB-wt) and LinB-H272F. (A) Stereo 2Fo-Fc electron density map contoured at 2σ for L9 and L16 loops in LinB-H272F (grey mesh). Protein residues shown as blue sticks. (B) Cartoon representations of LinB-wt (green; PDB ID: 1MJ5) and LinB-H272F (blue) structures and their superposition. Red dashed arrows depict the conformational changes observed in L9 and L16 loops. (C) Close-up views of the active site in LinB-wt (green) and LinB-H272F (blue) structures, and their superposition. Note that the presence of bulky and aromatic phenylalanine (F272) disturbs the positioning of the remaining catalytic triad residues (D108 and E132). Hydrogen bonds are shown as dashed yellow lines.

The major dissimilarities between LinB-H272F and other LinB structures lie in the different active site conformations surrounding the L9 and L16 loops (**Figure 3A**). The L16 loop, connecting the β7 strand and the α9 helix, adopts an atypical conformation that allows it to protrude toward the α4 helix. Notably, this relocation allows the side chain of residue L31 at the tip of the L16 loop to shield the side chain of introduced F272, making non-polar hydrophobic contacts. Concomitantly, the α4 helix and its adjacent L9 loop are rearranged too. Specifically, the α4 helix extends by a half turn on its proximal end while the L9 loop moves to shield the L16 loop, keeping it in its flipped-in conformation. The overall analysis of the LinB-H272F structure reveals major backbone changes that distinguish this structure from other LinB structures determined up to now.

### The bulky phenylalanine disturbs the catalytic triad

The catalytic machinery of all HLDs relies on the conserved aspartate-histidine-glutamate (**Figure 3B**) or aspartate-histidine-aspartate catalytic triad.^14^ The crystal structure of LinB-H272F reveals the altered positioning of the catalytic acid (E132) and the nucleophile aspartate (D108). The bulkier side chain of phenylalanine placed in the middle of the catalytic triad may have altering effects for several reasons. Firstly, it is one atom larger than the imidazole ring of histidine, causing spatial stress on the neighboring residues. Secondly, due to its inability to make hydrogen bonds, the hydrogen-bond network of the catalytic triad is abolished.

The histidine (H272) canonically interacts with E132 through a hydrogen bond. Consequently, E132 forms an additional hydrogen bond with A247 of the L16 loop (**Figure 3C**). However, these interactions are lost in the LinB-H272F structure. Moreover, E132 is flipped-out, not anchoring the above-located L16 loop, which consequently adopts a new, atypical conformation. The effect of the phenylalanine mutation is also observed in the positioning of the nucleophile aspartate (D108), located on the opposite side of the catalytic triad. The proper positioning of D108 is key for the S_N_2 catalytic mechanism. In the LinB-H272F, the carboxylate group of D108 is rotated from its canonical position compared to the LinB-wt (**Figure 3C**).

### An enzyme active site pocket rerouting

Although often overlooked, channels and tunnels are essential parts of enzymes. Over 60% of enzymes have their active sites accessible through channels and tunnels.^30–31^ They are essential for the substrate specificity, entrance of solvent molecules, synchronization of catalytic steps and reactions with more substrates, and prevention of cellular damage by reactive metabolites.^32–33^ Mutations that change the access tunnels can often affect enzyme activity,^34–36^ specificity,^37–39^ stability,^13, 27, 40^ and enantioselectivity.^41–43^

All HLDs have enzyme access tunnels that connect the active site with the enzyme surface.^44^ Our previous research showed that the morphology of these tunnels, physicochemical properties, and dynamical features are essential for productive catalysis and substrate specificity.^23, 45–46^ So far, up to four different access tunnels have been described in the HLD family (dubbed p1 to p4).^47^ In the LinB-H272F, the rearrangement of the L9 and L16 loops caused a major rerouting, resulting in the opening of an as-yet-unobserved enzyme access tunnel, hereafter referred to as p5 tunnel (**Figures 4** and **S6**). The p5 tunnel is markedly voluminous, uniquely going through the relocated L16 loop in the LinB-H272F structure. Inspection of the electron density map unambiguously identified several ligand molecules, including the CAPS buffer (N-cyclohexyl-3-aminopropanesulfonic acid), Tris buffer (tris(hydroxymethyl)aminomethane), and a phosphate ion occupying the p5 tunnel (**Figure 4B-C**). All these ligands originated from the crystallization of the mother liquor. While the canonical p1 and p2 tunnels are substantially reduced, newly opened p5 is the dominant tunnel in the LinB-H272F structure.

**Figure 4:**
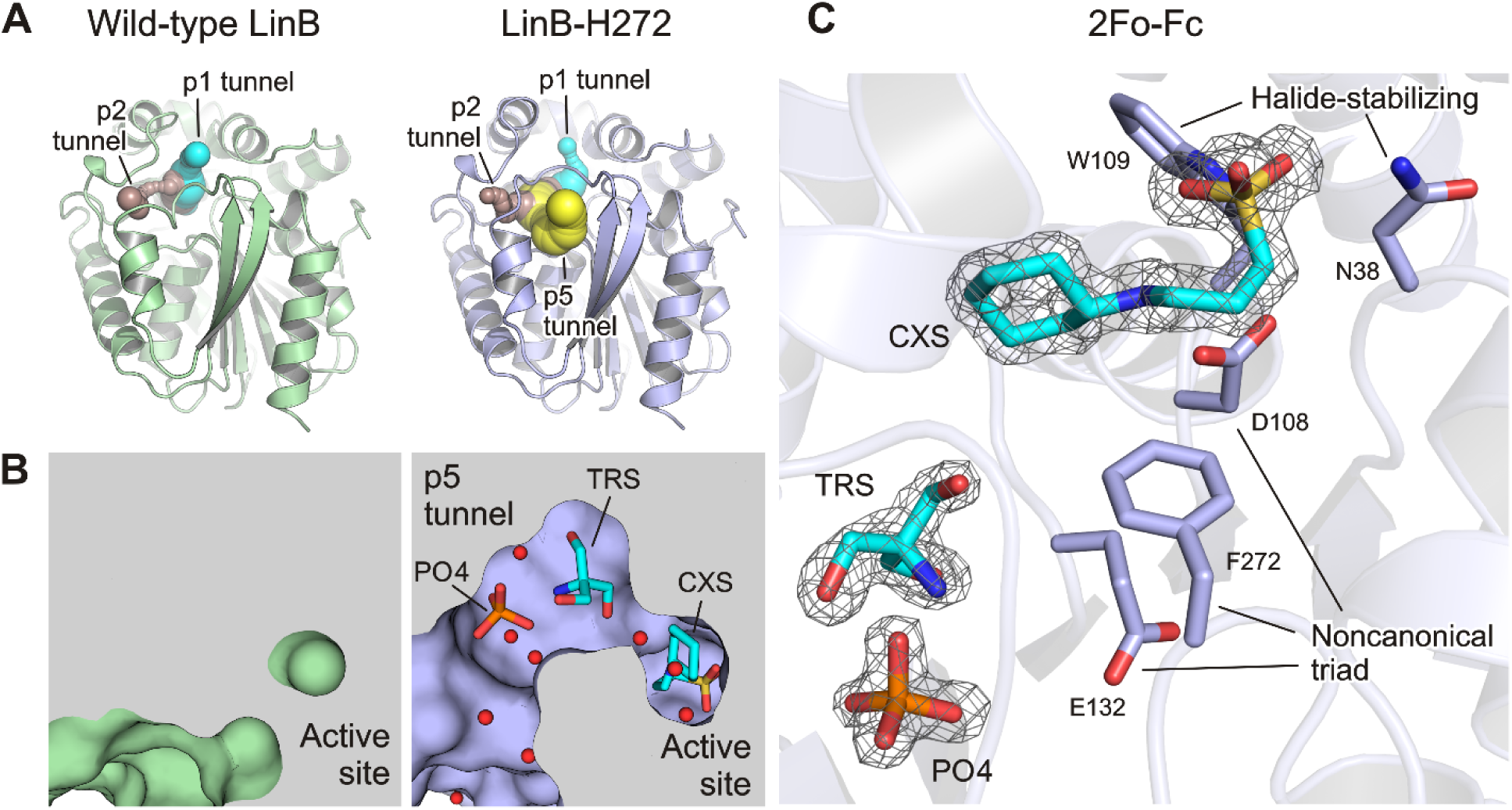
Rerouting of the active site pocket. (A) Visualization of enzyme access tunnels in LinB-wt (green; PDB ID: 1MJ5) and LinB-H272F (blue). Note that along with canonical p1 (cyan) and p2 (brown) tunnels, a newly open p5 tunnel (yellow) is exclusively present in LinB-H272F but not in LinB-wt. (B) Cutaway surface representations of LinB-wt (green) and LinB-H272F (blue) active site pocket. Note that the p5 tunnel in LinB-H272F is voluminous, capable of accommodating ligand molecules such as N-cyclohexyl-3-aminopropanesulfonic acid or CAPS (CXS), tris(hydroxymethyl)aminomethane or Tris buffer (TRS), and a phosphate ion (PO4). Water molecules are shown as red spheres. (C) 2Fo-Fc electron density (contour level 2σ) in the p5 tunnel of LinB-H272F structure. Catalytic pentad residues are depicted as blue sticks.

These crystallographic observations highlight that the histidine-to-phenylalanine mutation may affect the initial chemical step of the dehalogenation reaction (S_N_2) and as well as the binding and ligand transport through the enzyme access tunnel rerouting. As modelled in **Figure S7**, the replacement of the catalytic histidine base with an asparagine causes no observable spatial stress, and more importantly, it still supports the hydrogen bonding between the catalytic triad residues.

### Protein protonation states in neutral and alkaline pH

The crystals of LinB-H272F were obtained in an alkaline environment (pH 10.5). We, therefore, wondered whether the specific structural changes observed in the LinB-H272F could be a crystallographic artefact. To address this concern, we performed molecular dynamics (MD) simulations with LinB-wt, LinB-H272F, and LinB-H272N at neutral (pH 7.5) and alkaline environments (pH 10.5).

In the case of the H272N mutant, whose atomic structure has not been reported yet, a Rosetta model based on the *in silico* mutation of LinB-wt was used instead. The hydrogen atoms of LinB variants were predicted for pH 7.5 and 10.5 employing the H++ server.^48^ According to the calculations, the p*K*_a_ value of the catalytic histidine (H272) is 10.0, resulting in a doubly protonated histidine at pH 7.5 (**Figure 5A**). Increasing pH to 10.5 leads to a singly protonated histidine as a consequence of the N_δ_ deprotonation (**Figure 5B**). This difference in histidine protonation has a wide impact. Namely, at pH 7.5, the doubly protonated histidine of LinB-wt participates in an extended hydrogen bond network, which involves: (*i*) the catalytic residues D108 and E132 located in the loops L7 and L9, respectively, and (*ii*) the backbone nitrogen atoms of residues A247 and L248 located in the loop L16. Catalytic E132, as a part of loop L9, represents a link between H272 and loop L16. However, this connection is lost at pH 10.5 due to the loss of the proton at N_δ_. Hence, the flexibility of the loops L9 and L16 can increase.

**Figure 5:**
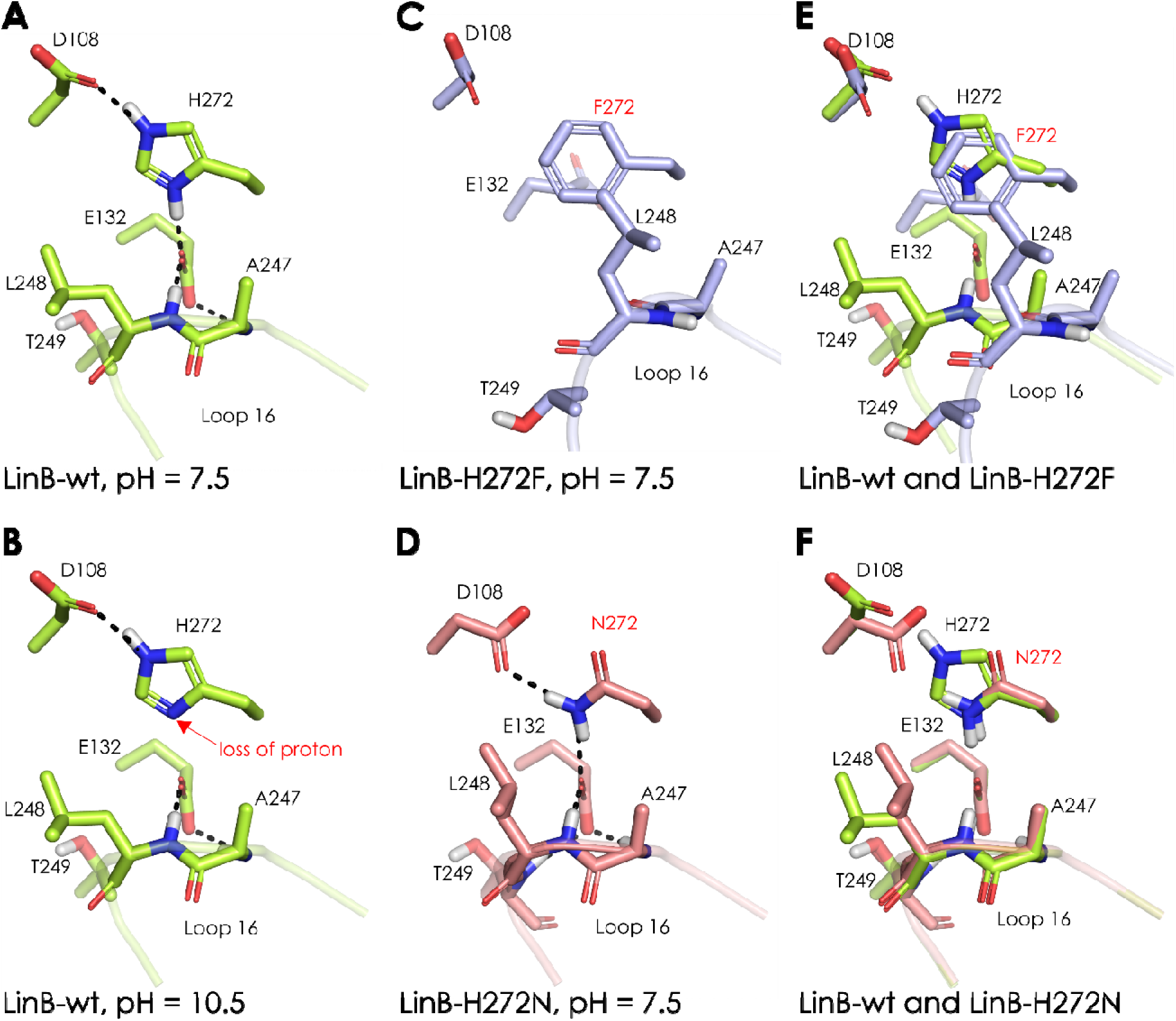
Protonation states of the LinB variants at different pH values and the hydrogen-bond network. (A) LinB-wt at pH 7.5. (B) LinB-wt at pH 10.5. At pH 10.5, the N_δ_ atom of H272 is deprotonated. (C) LinB-H272F at pH 7.5. (D) LinB-H272N at pH 7.5. (E) Superimposition of LinB-wt with LinB-H272F. (F) Superimposition of LinB-wt and LinB-H272N. For simplicity, only loop L16 is represented as a cartoon. The dashed black lines depict hydrogen bonds.

The mutation of H272 to F272 causes disruption of the whole hydrogen-bond network described above. As a result, the loops L9 and L16 are not in a firm position anymore (**Figure 5C**). This could be the reason why the catalytic residues adopt new conformations. In contrast to the LinB-H272F mutant, the Rosetta model of the H272N mutation showed that most of the interactions are preserved (**Figure 5D**). Only a small change in the D108 orientation occurs after 10 ns of the MD simulation, which causes the loss of the hydrogen bond with N272 (**Figure S8**). Superimpositions of H272F and H272N mutants with LinB-wt (**Figure 5E and F**, respectively) show that the conformation of the residues in the studied region of the H272F mutant changed substantially, while only minor changes can be observed for the H272N mutant.

### Protein dynamics and enzyme access tunnels

MD simulations were performed with the LinB variants at pH values 7.5 and 10.5, as described in the previous subsection. The simulations ran in duplicates for a total aggregated time of 1 μs, during which the systems remained stable according to the RMSD time variation (**Figure S9**). The flexibility of the proteins was assessed by the B-factors of the backbone atoms (**Figure 6**), for which the following observations were made: (*i*) LinB-wt and LinB-H272N at pH 7.5 have similar flexibilities (except for α5’ that is higher in the case of H272N), and they are the least flexible of all the tested systems. (*ii*) At pH 10.5, the flexibility of LinB-wt increases, especially in the regions around residues 159 (the beginning of helix α5’), 145 (loop L9), and 260 (loop L16). The B-factor of the loop L16 increases to the value found in LinB-H272F. (*iii*) LinB-H272F at both pH values has an increased flexibility of the loops L9, L14, and L16 compared to LinB-wt and LinB-H272N at pH 7.5. Interestingly, even in the MDs of the LinB-H272F Rosetta model (i.e., a model with an initial conformation identical to that of LinB-wt; **Figure S10**), the flexibility of loops L9, L14, and L16, and helix α5’ at pH 7.5 is increased compared to LinB-wt at the same pH, and the flexibility is comparable to the MDs with the crystal structure of LinB-H272F.

**Figure 6:**
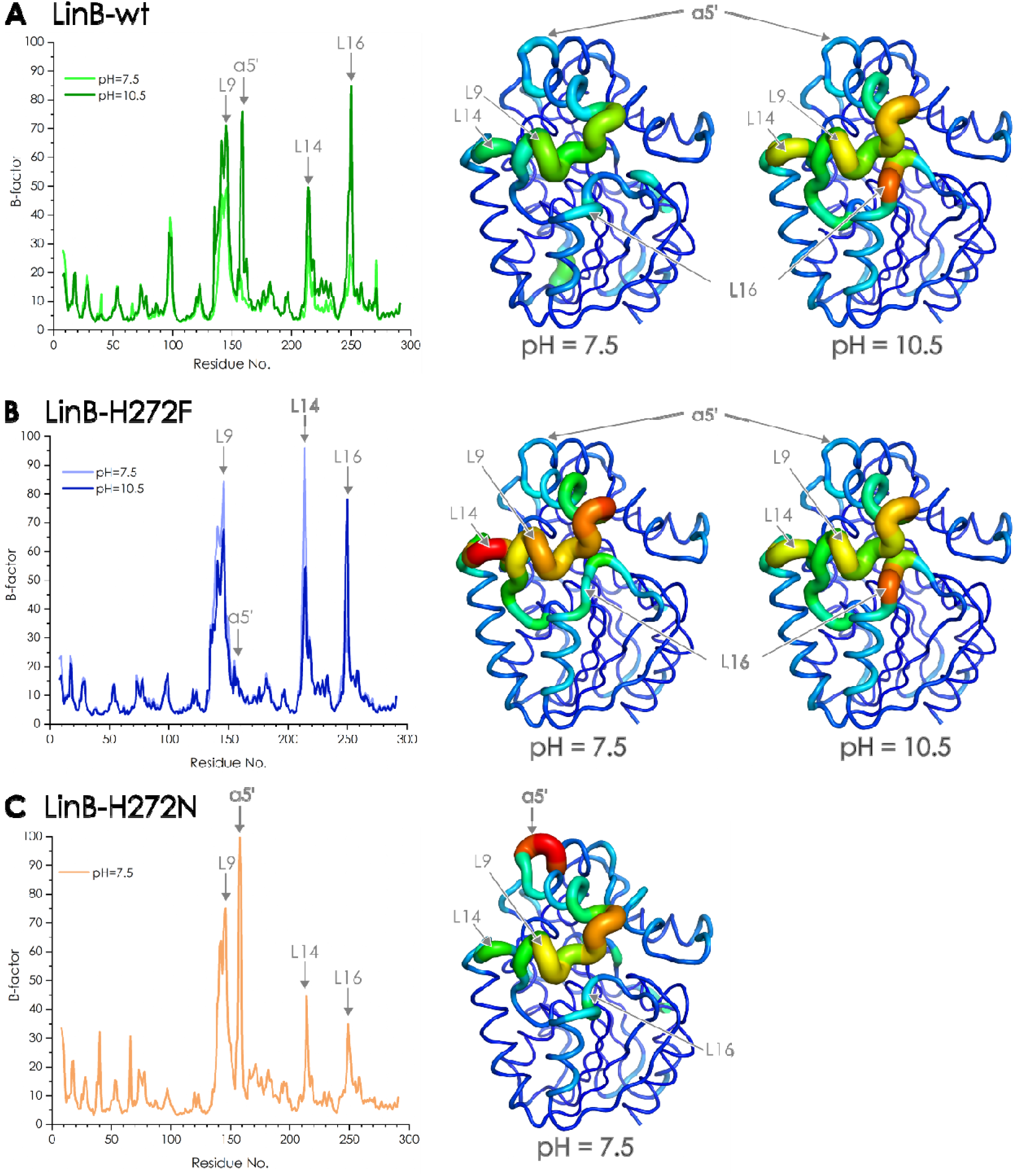
Protein backbone flexibility quantified by B-factor values from MD simulations. (A) LinB-wt, (B) LinB-H272F (B), and (C) LinB-H272N. The results, calculated from the respective MD simulations, are shown for all three proteins at pH values 7.5 and 10.5 (in the case of LinB-H272N only pH 7.5), depicted as graphs and as putty tube representations. A hotter (redder) and thicker putty tube indicates a higher B-factor (higher flexibility); the regions with more pronounced flexibility changes are indicated.

The observed differences in the flexibilities can be explained by the hydrogen-bond network involving the catalytic residues (D108, E132, and H272) and the residues of the loop L16 (A247, L248, and T249), as summarized in the previous subsection and **Figure 5**. To verify those observations in a dynamic context, we analyzed several interatomic distances inside the hydrogen-bond network during the MD simulations (**Figure S11**). LinB-wt and LinB-H272N at pH 7.5 conserve those interactions for most of the simulations (except for D108-N272; see **Figure S8**). However, at pH 10.5 LinB-wt loses the H-bond network for most of the MDs, conserving only the contact between D108 and H272. Similarly, LinB-H272F loses all the contacts inside the mentioned hydrogen-bond network. This supports the hypothesis that the disrupted hydrogen-bond network is the reason for the higher flexibility of the loops L9 and L16 and the consequent appearance of the p5 tunnel.

The differences in the values of kinetic rates obtained by transient kinetic experiments (**Figure 2**) can be caused by the orientation and flexibility of the catalytic D108, one of the key catalytic triad residues in this step. The computed dihedral angle of the catalytic D108 side chain in wild-type and the LinB-H272N mutant (**Figure S12A**) shows similar distributions (peak at 64.0° and 64.4°, respectively), while in the LinB-H272F mutant, it is shifted (**Figure S12B**, distribution peak at 59.0°). Despite the small differences, this might lead to different reactivity in the LinB-H272F mutant.

The tunnels in the crystal structures and MDs were computed and analyzed by CAVER 3.02.^49^ In the crystal structure of LinB-H272F, three important tunnels (p1, p2, and p5) were found in contrast to LinB-wt, with only two tunnels (p1 and p2). **Table 1** shows the properties of the tunnels for: (*i*) crystal structures of LinB-wt and LinB-H272F (**Figure 4A**), and (*ii*) MD simulations (LinB-wt, LinB-H272F, and LinB-H272N) at pH 7.5 and 10.5 (**Figure S13**). The p5 tunnel of the LinB-H272F crystal is relatively wide (bottleneck radius of 1.56 Å) and it has potentially high importance in the transport of small substances (solvent, ions, substrates), ranking as the best tunnel (priority of 0.673) according to CAVER. The remaining tunnels, p1 and p2, changed their shapes and radii slightly compared to the LinB-wt but not remarkably. The results for the MD simulations provided additional information in terms of the protein behaviour in solution. Namely, the relevant tunnels observed in LinB-wt and LinB-H272N at pH 7.5 were mainly p1 and p2, while p5 was not found in either of them. In contrast, the newly found tunnel, p5, is present in LinB-H272F at both neutral and alkaline pH. Although it was not the preferred tunnel (p1 always ranked first), p5 was detected in almost half of the snapshots, and it displayed the maximum bottleneck radius of 1.74 Å, which means that it may be relevant for the transport of water and ligand molecules. Moreover, LinB-wt displays the new p5 tunnel at pH 10.5, driven by the increased flexibility of loop L16 (**Figure 6A**), unlike what was found for the same protein at pH 7.5. The p5 tunnel in LinB-wt at pH 10.5, although present only in 15% of the snapshots, displayed a maximum bottleneck radius of 1.64 Å, sufficient for transporting different molecules. In the MDs of the LinB-H272F Rosetta model (i.e., with an initial conformation identical to that of LinB-wt) at pH 7.5, the main tunnels were p1 and p2, but p5 tunnel could already be detected, although in very few frames (3.8%) and with small priority (**Figure S13F**). This suggests that p5 is not a consequence of the initial crystallographic structure, but it very likely occur in solution.

**Table 1:**
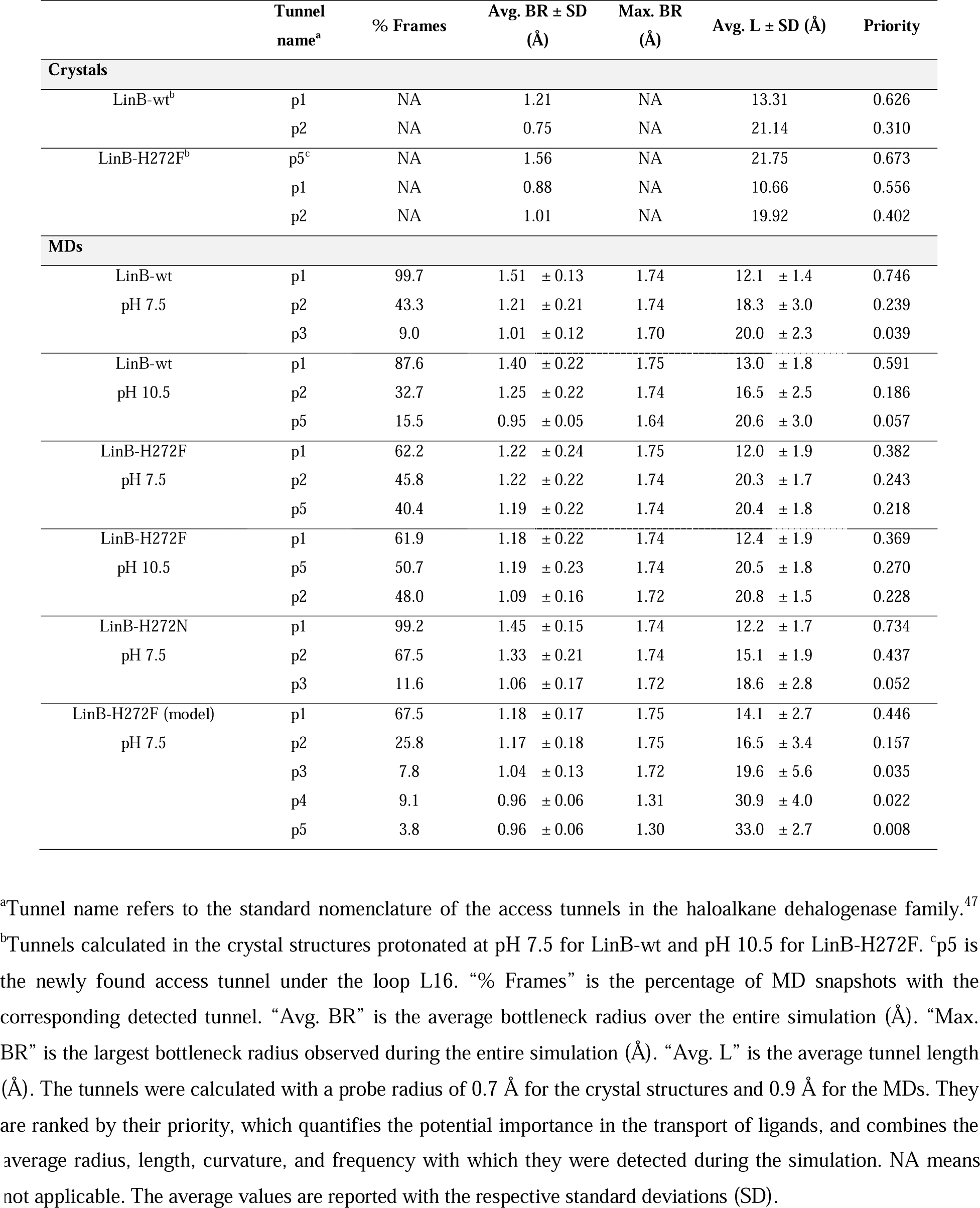
Properties of the tunnels ordered by priority calculated in the crystal structures and MD simulations of different LinB variants at different pH values.

### Ligand transport via tunnels

The tunnels calculated for the crystal structures and MD simulations were examined in terms of their importance for the transport of ligands. The investigated ligands were the same as in the case of transient kinetic experiments (**Figure 2**), where Cl^-^ ion was considered as a product and 1-chlorohexane as a substrate. The binding of the substrate and unbinding of the product were studied by adaptive steered molecular dynamics (ASMD) simulations at pH 7.5 for all three LinB variants (LinB-wt, LinB-H272F, and LinB-H272N). We estimated preferences of these ligands for the tunnels p1, p2, and p5. The ASMD method applies a constant external force on two atoms that can be either pulled together or pushed away from each other to simulate the binding or unbinding of ligands, respectively. Since the p5 tunnel is not naturally present in either LinB-wt or LinB-H272N at pH 7.5, we found the corresponding tunnels with a smaller probe (0.7 Å) to set the direction for the (un)binding (**Figure S14**). However, due to the different conformations of the loop L16, the identified p5 tunnels have very different geometries compared to LinB-H272F.

The binding of 1-chlorohexane (substrate) was simulated through all three the aforementioned tunnels (**Figure 7**, left panels). In the case of LinB-wt, the substrate goes smoothly through the p1 tunnel. When attempting to enter through tunnels p2 and p5, the substrate uses p1 instead in the end, revealing that the transport through the p2 and p5 tunnels is highly unfavorable. Moreover, the potential of mean force (PMF) is lower for the p5 tunnel, suggesting that this is a preferred binding path. In the LinB-H272N mutant, the substrate tries to enter through p5 but eventually slides over the protein surface and enters through p1. In this variant, the ligand can enter successfully through p2 as well, although with a higher PMF (less favorable). In the energy profiles of all three proteins (**Figure 7**, left panels), we see a clear preference for p1, except for LinB-H272F, where the substrate successfully goes through p5 as well.

**Figure 7:**
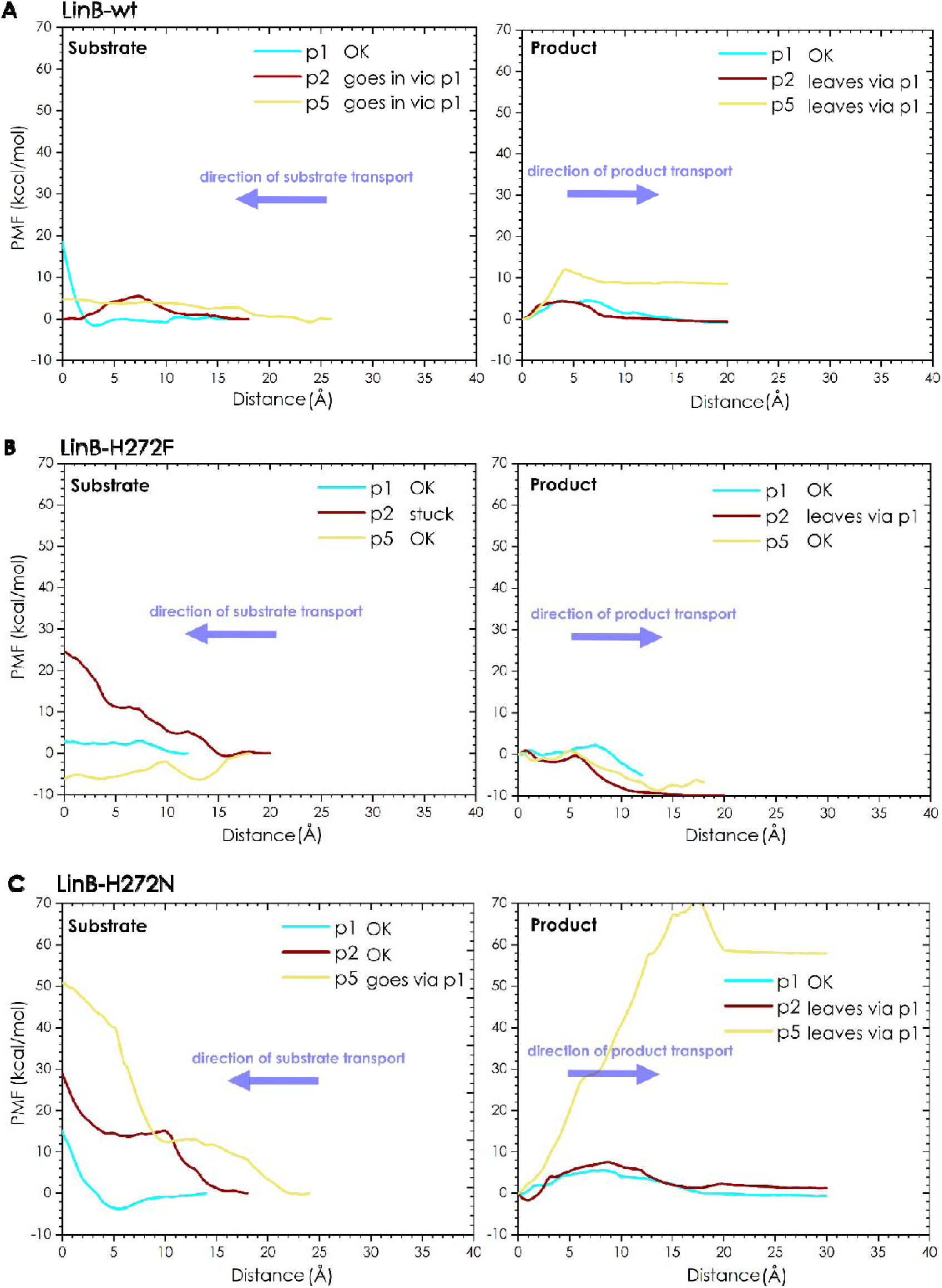
Potential Mean Force (PMF) profiles from ASMD simulations. Simulations with 1-chlorohexane (substrate) are presented on the left and with Cl^-^ ion (product) on the right, for LinB-wt (A), LinB-H272F (B), an LinB-H272N (C). Blue arrows depict the transport direction for the substrate and the product. The distance value of 0 represents the starting/finishing point of the ligand transport. An inset of each panel summarizes the final result of the simulation for all three tunnels.

The unbinding of Cl^-^ ion (product) was simulated through the same tunnels as for the substrate (**Figure 7**, right panels). In LinB-H272F, the ion can leave through the tunnels p1 and p5, with similar PMF profiles. In the case of tunnel p2, the product cannot pass, and finally, it leaves through p1. In LinB-wt and LinB-H272N, the ion can leave only through the tunnel p1. In the case of the other two tunnels, the ion tries to leave through the given tunnel, but after being pushed for a certain distance, it leaves through p1 instead. The energy profiles (**Figure 7**, right panels) show that when the ion is pushed through the non-obstructed tunnels, the unbinding process has a very low barrier, and the transport of the ion is not hindered. However, in LinB-H272F additional preferred transport via the tunnel p5 appears and might be even higher than for the tunnel p1.

## Discussion

The HaloTag is a widely used protein labelling in live cells, fixed samples, or even *in vitro*, with the flexibility to swap ligands for imaging, tracking, or functional studies.^16, 18, 50^ It is an engineered protein labelling system derived from bacterial haloalkane dehalogenase DhaA, in which the catalytic histidine is mutated to block the hydrolysis step of the native reaction.^17^ In the wild-type enzyme, this histidine activates water to cleave a covalent intermediate with a haloalkane substrate, but in HaloTag, the mutation makes the reaction irreversible. In this work, we combined kinetic, structural, and *in silico* analyses to obtain this understanding and explain why there are differences in the performance of the two mutants (H272N and H272F).

Both LinB mutants of the catalytic histidine (H272F and H272N) block the hydrolytic step in the HLD reaction. However, only the H272N variant can form a stable covalent bond at the S_N_2 step which is irreversible. On the other hand, the H272F mutant can reverse the S_N_2 step and at this stage it can release a Cl^-^ ion from the active site, a feature not seen in other native HLDs. This suggests a possible application of H272F variants of HLDs in transhalogenation reactions^24^ with milder reaction conditions compared to the conventional synthetic approach. Consequently, this can provide a green and sustainable way with the possibility of recycling the halogenated compounds. However, for industrial applications, enzymatic properties need to be tailored to obtain good solubility, stability, and enzymatic activity.^50^ This can be successfully achieved by rational design, for which the structural information is crucial.^23^ To this end, to understand why there are differences in the performance of the two mutants (H272N and H272F), we combined a study of enzymatic activity with structural studies.

Our transient kinetic experiments revealed the dependence of the S_N_2 step on the mutation type and pH value. At neutral pH, LinB-H272F and LinB-H272N mutants exhibit 1000- and 10-fold lower rates compared to the LinB-wt, respectively. The S_N_2 step was nearly irreversible for the LinB-H272N mutant but highly reversible for the LinB-H272F mutant. On the other hand, the alkaline pH for LinB-H272F mutant affected the activity in the way that Cl^-^ product was released at the level of the intermediate (during the S_N_2 step), which highlights the importance of this mutation in transhalogenation reactions.^24^ To understand what contributes to the different enzymatic performances of these two mutants, we solved the structure of LinB-H272F with a resolution of 1.55 Å. It revealed differences in the conformation of loops L9 and L16 compared to LinB-wt, and a wide access tunnel (p5), never seen in the HLD family before. A TRIS molecule, a phosphate anion, and a CAPS molecule were found in the tunnel. The size of these molecules was that of a typical size of the HLD substrates.^23^ For a closer examination of this phenomenon, MD simulations were performed in neutral and alkaline environments. The reason for the different conformation of loops L9 and L16 was the disruption of an extended hydrogen-bond network involving the catalytic histidine and residues in loops L9 and L16. Therefore, the disruption was caused by the H272F mutation, but not the H272N mutation, or by the deprotonation of the catalytic histidine in LinB-wt due to the high pH value. This disruption was followed by the appearance of a new tunnel, p5, which was not found in the LinB-H272N mutant. ASMD simulations showed that the substrate (1-chlorohexane) and the product (Cl^-^ anion) preferred the p1 tunnel in all three LinB variants, together with the p5 tunnel in the LinB-H272F mutant. This confirmed the observations from the X-ray experiments that the newly found p5 tunnel in the LinB-H272F mutant plays a role in the substrate/product transport. Creation of a completely new access tunnel in proteins with a buried active site is extremely challenging.^34^

The power of a single-point mutation as a tool for creating new enzymatic activities is not limited only to the HLD family.^51^ Recently, Tripathi and Dutta Dubey^9^ reported that a mutant of homoserine kinase gained a secondary catalytic activity as an ATPase. In another study, a long-distance effect of the single-point mutation was demonstrated in AEE epimerase.^11^ The mutation far from the capping loop affected the conformation in that region, causing the accommodation of a non-native substrate. The substrate promiscuity and improved native reactivity, important for industrial applications, were achieved in a single-point mutant of cytochrome P450GcoA.^10^ This work highlights that elucidating structure–function relationships is essential for guiding rational design strategies, ultimately enabling the creation of enzymes with enhanced or novel catalytic functions.

## Conclusions

We investigated the impact of a single-point mutations in the catalytic histidine on the structure and function of the haloalkane dehalogenase LinB. We demonstrated that S_N_2 chemical step depends on the mutation type. Namely, H272N mutant forms a strong covalent bond in an irreversible reaction S_N_2 step. On the other hand, the S_N_2 step becomes reversible in the case of the H272F mutant, while releasing the Cl^-^ ion at this stage. This feature is important in transhalogenation reactions. To understand what contributes to the different activity performances of these two mutants, we solved the X-ray structure of the LinB-H272F mutant, where we observed a wide access tunnel (p5), never seen in the HLD family before. MD simulations showed the disruption of an extended hydrogen-bond network linking the catalytic histidine and loops L9 and L16, far from the active site. This rearrangement caused the opening of the new p5 tunnel in the H272F variant but not in H272N. Together, these findings underscore how a single, precisely chosen amino acid substitution can reprogram enzyme reactivity, alter reaction reversibility, and create entirely new structural features — offering opportunities for engineering novel enzymatic functions.

## Materials and methods

### Site-directed DNA mutagenesis and molecular cloning

Standard site-directed mutagenesis generated the LinB-H272F and LinB-H272N mutants. Briefly, the QuikChange™ Site Directed Mutagenesis Kit (Stratagene, Santa Clara, CA, USA) was used for mutagenesis. Specific forward and reverse primers encoding the mutation were designed and used in PCR-based amplification. The resulting PCR products were incubated with DpnI restriction endonuclease to remove the template plasmids and then transformed into *E. coli* chemo-competent cells via a heat-shock method. The error-free clones were confirmed by DNA sequencing. The genes of LinB mutants were cloned into pAQN expression plasmid between EcoRI and HindIII sites.

### Protein expression and purification

The expression plasmids were transformed into *E. coli* BL21(DE3) cells which were plated on agar plates with ampicillin (100 µg·mL^−1^) and grown overnight at 37°C. Then, the cells were cultured in LB medium at 37°C until OD_600_ of ∼0.6. After the cultivation, the expression of the protein was induced by the addition of isopropyl-β-D-thiogalactopyranoside (IPTG) to a final concentration of 0.2 mM. The cells were incubated at 20°C overnight and then harvested by centrifugation. The pellet was resuspended in a purification buffer (20 mM potassium phosphate buffer with pH 7.5, 500 mM NaCl, 10 mM imidazole), the cells were disrupted by sonication using the Ultrasonic Processor UP200S (Hielscher, Germany), and centrifugated (20,000×g, 1h, 4°C). After centrifugation, the supernatant was collected, and the His-tagged enzyme was purified using Ni-NTA Sepharose column HR 16/10 (Qiagen, Germany). The enzyme was eluted with 10 and 60% of the elution buffer (20 mM potassium phosphate buffer with pH 7.5, 500 mM NaCl, 500 mM imidazole) and dialyzed in 2 L of 50 mM potassium phosphate buffer, pH 7.5. The sample was then further purified using gel filtration on Äkta Purifier FPLC (GE Healthcare, Sweden) equipped with HiLoad 16/600 Superdex 200 pg column (Cytiva, Sweden). The purified protein was concentrated to the final concentrations using Centrifugal Filter Units AmiconR Ultra-15 UltracelR-10K (Merck Millipore Ltd., Ireland) equilibrated with the gel filtration buffer.

### Transient kinetic experiments

The transient kinetic experiments were performed at 27°C (300 K) in 100 mM glycine solution with pH 7.5 or 10.5, depending on the collected dataset. The components were rapidly mixed using the Stopped-Flow instrument SFM 3000 combined with the MOS-500 spectrometer (BioLogic, France). The reaction was initiated by rapid mixing 75 μL of an enzyme (LinB-wt, LinB-H272F, or LinB-H272N) with 75 μL of sodium chloride or 1-chlorohexane at various concentrations. Kinetic data were collected by monitoring the change of native tryptophan fluorescence (295 nm excitation, 320 nm cut-off filter emission). The change of native tryptophan fluorescence was forged by the binding-unbinding of Cl^-^ to catalytic W109. After mixing, the resulting concentrations of enzymes were fixed at 3.5–4.2 μM while the resulting concentration of sodium chloride and 1-chlorohexane ranged from 0 to 2000 mM and 0 to 1860 μM, respectively. Each kinetic trace was measured in seven consecutive replicates and averaged.

### Transient kinetics data analysis

In the case of the chloride anion (product) binding, where the rapid binding within the instrument dead time (0.3 ms) occurred, the level of equilibrium fluorescence was normalized and plotted towards the chloride ion concentration. The concentration dependence was fitted with a hyperbolic function (**Equation 1**), allowing us to determine the value of the dissociation constant, *K*_d_.

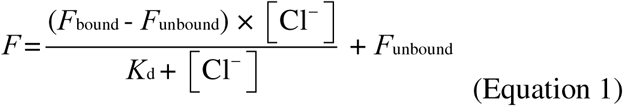

Kinetic traces, collected upon mixing an enzyme with 1-chlorohexane (substrate) were fitted by using a single exponential equation (**Equation 2**), and the values of obtained exponential observed rates (*k*_obs_) were plotted against the 1-chlorohexane concentration determined by gas chromatography. The concentration dependence of the observed rate was fitted by the hyperbolic function (**Equation 3**), derived from a two-step mechanism (**Equation 4**) consisting of substrate binding, followed by a chemical step.^52^

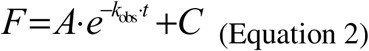

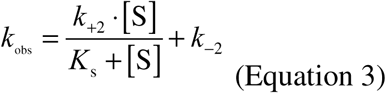

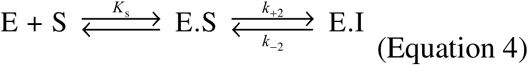

In the case of the wild type, the intercept of the hyperbolic function (**Equation 4**) is not *k*_-2_ solely, but rather the sum of *k*_-2_ and *k_+_*_3_, as the reaction proceeds to subsequent catalytic steps. By employing a simplifying assumption that the *k*_cat_ value for LinB-wt converges towards *k_+_*_3_ (following rate-limiting hydrolytic step),^53^ the *k*_-2_ value for the wild type can be obtained by subtracting *k*_cat_ from the intercept.

Due to the very slow 1-chlorohexane processing by LinB-H272N at pH 10.5 and no detectable exponential fluorescence decrease within the measured time window (100 seconds), only the upper limits of the chemical step rate constants *k*_+2_ and *k*_-2_ (0.01 s^-1^) were estimated. In this case, the substrate binding dissociation constant value, *K*_s_, was derived based on the concentration dependence analysis of the initial fluorescence quench caused by the substrate binding. The dependence was fitted using the hyperbolic function (**Equation 1**) as in the case of rapid-equilibrium binding of chloride anions. In the case of substrate binding by LinB-H272F at pH 10.5, where a more complex behavior of the enzyme was detected (**Figure S4**), numerical data fitting was performed using KinTek Explorer 10 (KinTek Corporation, USA) to account for the sign of conformational selection substrate binding mechanism and chloride release after the alkyl-enzyme intermediate formation (**Equation 5**). Numerical integration of the rate equations, searching a set of kinetic parameters that produce a minimum χ^2^ value, was performed using the Bulirsch–Stoer algorithm with adaptive step size, and nonlinear regression to fit data was based on the Levenberg–Marquardt method. Residuals were normalized by sigma value for each data point. The standard error was calculated from the covariance matrix during nonlinear regression. The final values of rate constants were used to derive the parameters of substrate binding dissociation constant (*K*_s_) and rate constants of the alkyl-enzyme intermediate formation (*k*_+2_) and breakup (*k*_-2_) to quickly compare the values.

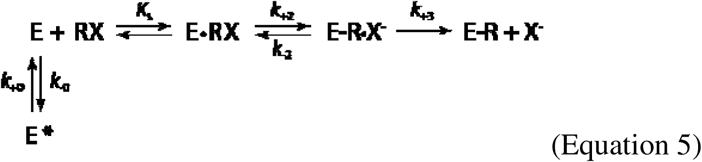

### Crystallization and diffraction data collection

The LinB-H272F protein was concentrated to 10 mg·mL^-1^. The crystallization was performed by using the hanging-drop vapor-diffusion method at 20°C. LinB-H272F crystals were obtained in a crystallization buffer containing 1.2 M sodium di-hydrogen phosphate, 800 mM di-potassium hydrogen phosphate, 200 mM lithium sulfate, and 100 mM CAPS (pH 10.5). The drops were formed by mixing 1 µL of protein solution and 1 µL of the reservoir solution, and they were equilibrated against 500 µL of the reservoir solution. The crystal growth was monitored by stereomicroscope Olympus-SZX10 (Olympus, Czech Republic). The obtained crystals were soaked into the crystallization solution supplemented with 20% glycerol, and flash-frozen in liquid nitrogen. The diffraction data were collected at 100 K at PXIII beamline at the Swiss Light Source (SLS), using a Pilatus 2M-F detector, multi-axis PRIGo goniometer and at a wavelength of 0.999 Å.

### Crystal structure determination and refinement

The diffraction data were processed by XDS,^54^ scaled, and merged in Aimless.^55^ The crystal structure was solved by a molecular replacement using Phaser^56^ implemented in Phenix package,^57^ and employing the stricture of wild type LinB (LinB-wt; PDB ID: 1MJ5), as the search model. Structural refinements were carried out using phenix.refine,^57^ alternating with multiple cycles of manual model building in Coot.^58^ The structural representations were prepared by PyMOL.^59^ The structural coordinates and structure factors files were deposited to the Protein Data Bank^60^ under the PDB ID code: 7NFZ.

### Molecular dynamics simulations and analysis

The 3D structure of LinB-H272F (PDB ID: 7NFZ; this work) and LinB-wt (PDB ID: 1MJ5^61^) were used as input. The solvent, co-crystallization molecules, and ions were removed, and the double side chains were corrected to keep only the most populated conformations using the *pdb4amber* module of AmberTools 16.^62^. The hydrogen atoms were predicted using the H++ server,^48^ calculated in an implicit solvent at both pH 7.5 and 10.5, 0.1 M salinity, an internal dielectric constant of 10, and external of 80. The original crystallization solvent was added and the *tLEaP* program of AmberTools 16 was used to prepare the topology and coordinates files. For that, the force field ff14SB^63^ was specified, Na^+^ and Cl^-^ ions were added to neutralize the system and achieve 0.10 M concentration of NaCl salt, and a truncated octahedral box of TIP3P waters^64^ with the edges at least 10 Å away from the protein atoms was added.

The molecular dynamics (MD) simulations were carried out with PMEMD.CUDA^65–66^ module of AMBER 16.^67^ In total, five minimization steps and twelve steps of equilibration dynamics were performed prior to the production MD. The first four minimization steps, composed of 2,500 cycles of the steepest descent algorithm followed by 7,500 cycles of conjugate gradient, were performed as follows: (*i*) in the first one, all the atoms of the protein and ligand were restrained with a 500 kcal·mol^-1^·Å^2^ harmonic force constant; (*ii*) in the following ones, only the backbone atoms of the protein and heavy atoms of the ligand were restrained, respectively, with 500, 125, and 25 kcal·mol^-1^·Å^2^ force constant. A fifth minimization step, composed of 5,000 cycles of the steepest descent and 15,000 cycles of conjugate gradient, was performed without any restraints. The subsequent MD simulations employed periodic boundary conditions, the particle mesh Ewald method for treatment of the long-range interactions beyond the 10 Å cutoff,^68^ the SHAKE algorithm^69^ to constrain the bonds involving the hydrogen atoms, the Berendsen barostat^70^ at 1 bar, the Langevin thermostat with collision frequency 1.0 ps^-1^, and a time step of 2 fs. Equilibration dynamics were performed in twelve steps: (*i*) 20 ps of gradual heating from 0 to 310 K, under constant volume, restraining the protein atoms and ligand with 200 kcal·mol^-1^·Å^2^ harmonic force constant; (*ii*) ten MDs of 400 ps each, at constant pressure (1 bar) and constant temperature (310 K), with gradually decreasing the restraints on the backbone atoms of the protein and heavy atoms of the ligand with harmonic force constants of 150, 100, 75, 50, 25, 15, 10, 5, 1, and 0.5 kcal·mol^-1^·Å^2^; (*iii*) 400 ps of unrestrained MD at the same conditions as the previous restrained MDs. The energy and coordinates were saved every 10 ps. The production MDs were run for 500 ns using the same settings employed in the last equilibration step and performed in duplicate for each system. The two simulations of each type were combined into a single one using the *cpptraj*^71^ module of AmberTools 16, and aligned to the respective crystal structures by minimizing the root-mean-square deviation (RMSD) of the backbone atoms, excluding the very flexible terminal residues of each chain (6 terminal residues). The trajectories were analyzed using *cpptraj* and visualized using PyMol 2.3.2^59^ and VMD 1.9.1.^72^

### *In silico* mutagenesis

The structural models of LinB-H272F and LinB-H272N were constructed by *in silico* mutagenesis performed on the crystal structure of the LinB-wt, obtained from the RCSB Protein Data Bank^60^ (PDB ID 1MJ5). In the case of LinB-H272F, this approach was used only to verify some of the MD results gained based on the respective crystal structure. However, for LinB-H272N no 3D structure was available, hence the need to model it *in silico*. The crystallographic water molecules and ions were removed, and the protein chain was renumbered to start from position 1. The resulting structure was minimized by Rosetta using *minimize_with_cst* module. The minimization was performed according to Kellog and coworkers^73^. Both the backbone and the side chain optimization were enabled (*sc_min_only false*), the distance for the full atom pair potentials was set to 9 Å (*fa_max_dis* 9.0), the standard weights for the individual terms in the energy function were used with the constraint weight 1 (*constraint_weight* 1.0). The script convert_to_cst_file used the output from the minimization step for the creation of the cst file. To calculate the most stable conformers of each mutant, Protocol 16 was followed. For that, the *ddg_monomer* module of Rosetta was used according to Kellog *et al.*,^73^ incorporating the backbone flexibility. The soft-repulsive design energy function (*soft_rep_design weights*) was used for the side chains repacking and the backbone minimization (*sc_min_only false*). The optimization was performed on the whole protein without distance restriction (*local_opt_only false*). The previously generated constraints file was used during the minimization (*min_cst true*) to impose a restraint of 0.5 Å on the Cα atoms. The optimization was performed in three rounds with increasing weight on the repulsive term (*ramp_repulsive true*). The structure with the lowest energy (*mean false, min true*) was selected from the 50-iteration cycle (iterations 50), and it was used as the final result to obtain the minimized model of the mutant. All calculations used *talaris*2014^74–75^ force field.

### Access tunnel calculations

CAVER 3.02^49^ was used to calculate and cluster the tunnels in the crystal structures and during the MD simulations of the LinB variants. The analysis of the crystal structures was performed after the hydrogen atoms were added with H++^48^ at the respective pH. The tunnels were calculated during the MDs for all the 10 ps-spaced snapshots (in a total of 100,000 per system). We specified a probe radius of 0.9 Å for the MDs and 0.7 Å for the crystal structures, a shell radius of 3 Å, and a shell depth of 4 Å. The starting point for the tunnel calculation was defined by the geometric center of the two carboxylic oxygen atoms of the catalytic D108 residue. The clustering was performed by the average-link hierarchical Murtagh algorithm, with a weighting coefficient of 1 and a clustering threshold of 3.5 Å. The approximate clustering was allowed only when the total number of tunnels was higher than 20,000 and performed using 20 training clusters.

### Preparation of the chloride- and 1-chlorohexane-bound enzyme complexes

The unbinding of the chloride ion and binding of 1-chlorohexane were calculated using adaptive steered molecular dynamics (ASMD). To prepare the chloride ion, we modeled the HCl molecule using Avogadro.^76^ The HCl was saved as a MOL2 file and processed into the PDBQT file for the docking by MGLtools.^77^ The hydrogen atom was then deleted from the PDBQT file, and the charge of the chloride changed to –0.9999. The receptor PDBQT file was prepared by MGLtools as well. The grid box for the docking was calculated to contain the active site cavity, and the exhaustiveness of the docking was set to 100. The ion was then docked using the AutoDock Vina engine^78^ implemented in CaverDock.^79^ The best-docked pose with the lowest binding energy was selected, extracted, and converted to the MOL2 file. The complex for the MD simulation was then prepared with the following steps. The original crystallization solvent and the docked pose of the ion were added, solvent molecules clashing with the position of the ion were removed and the tLEaP program of AmberTools 16^62^ was used to prepare the topology and coordinates files. For that, the force field ff14SB^63^ was defined, Na^+^ and Cl^-^ ions were added to neutralize the system and achieve 0.10 M concentration of NaCl salt, and a truncated octahedral box of TIP3P^64^ water molecules with the edges at least 10 Å away from the protein atoms was added.

The molecule of 1-chlorohexane was also modeled in Avogadro^76^, minimized with the MMFF94 forcefield,^80^ and then saved as PDB file. The RESP charges for 1-chlorohexane were calculated by the Gaussian09_E.01 in R. E. D. Server.^81^ The parameterized MOL2 file from R. E. D. Server was then converted into a PDBQT file for docking by MGL tools.^77^ The end states of the minimization of the previously prepared complexes with the chloride ion were used as the starting point to set up the complexes with 1-chlorohexane. The chloride ion and all the solvent molecules were removed.

Then we calculated tunnels in the clean protein structures by CAVER 3.02.^49^ The tunnel calculation settings were kept as default apart from the probe radius. We started with the probe radius of 0.7 Å, which had to be lowered to find tunnels corresponding to the position of the p5 tunnel in the LinB variants (**Figure S14**). The tunnels corresponding to p1, p2, and p5 for LinB variants were extracted. The following steps prepared the systems containing the ligand molecule at the tunnel mouth. We calculated the vector between the first and the last tunnel sphere for each of the extracted tunnels. We extended this vector by 3 Å and saved the endpoint coordinates. These coordinates were then used to set up the constrained docking performed by CaverDock. The grid-box was set to include the whole protein and the position of the ligand outside of the protein. The chlorine atom was constrained to the end point coordinates, and we ran the constrained docking with exhaustiveness 1. We selected the best-docked pose as the one where the linear tail of 1-chlorohexane would be pointing outward from the protein structure so that the whole molecule would be positioned to go inside the tunnel. The topology, solvation, and water box were prepared in the same way as for the complex with the chloride ion.

### Molecular dynamics simulations with ligands

By using ASMD, we calculated the unbinding (chloride ion) and binding (1-chlorohexane) trajectories. The systems were equilibrated as described above for the molecular dynamics simulations, using the temperature of 300 K instead. The ASMD method applies constant external force on two atoms in the simulated systems to either push two atoms from each other or pull them together to simulate the unbinding or binding of the ligands, respectively.

For our purpose, we used the default values for setting up ASMD, which were found in the tutorial for AMBER. Used parameters: 25 parallel simulations, 2 Å stages, velocity 10 Å·ns^-1^, and force 7.2 N. The rest of the MD settings were set as in the last equilibration step. The ligand atom for steering was the chlorine atom in both cases. The protein atom for the steering is different for each tunnel. In each case, we selected a Cα atom in a residue that is opposite to the tunnel opening so that the ligand could be pushed or pulled in this direction (**Table S3**).

## Acknowledgments

The authors acknowledge the support of the RECETOX Research Infrastructure (No. LM2023069) and CZECRIN (No. LM2023049), funded by the Ministry of Education, Youth and Sports (MEYS) of the Czech Republic. This project was also supported by the European Union’s Horizon 2020 Research and Innovation Programme under grant agreements CETOCOEN Excellence (No. 857560) and CLARA (No. 101136607). Additional funding was provided through the ESIF-MEYS Johannes Amos Comenius Programme under the CLARA project (No. CZ.02.01.01/00/23_029/0008437), co-financed by the European Union and MEYS, and Czech Science Foundation (GA22-09853S and GX25-17329X). The article reflects the author’s view, and the Agency and European Comission are not responsible for any use that may be made of the information it contains. The authors are thankful to Swiss Light Source (SLS) synchrotron members for using their beamline facilities and help during data collection. We acknowledge CF BIC of CIISB, Instruct-CZ Centre, supported by MEYS CR (LM2023042) and European Regional Development Fund-Project “Innovation of Czech Infrastructure for Integrative Structural Biology” (No. CZ.02.01.01/00/23_015/0008175).

## Author Contributions

**M.T.**, **S.M.M.**, **T.G.**, and **H.B.** contributed equally to this work. **H.B.**, **V.N.**, and **M.M.** prepared protein samples, carried out the crystallization screenings, and optimized crystallization hits. **H.B.** and **M. M.** collected diffraction data and solved the protein crystal structure. **M.T.** performed transient kinetic experiments and analysis. **S.M.M.** and **O.V.** performed MD and ASMD simulations, respectively. **J.D.**, **D.B.**, **Z.P.**, and **M.M.** designed the project, supervised research, and interpreted data. **M.T.**, **S.M.M.**, **T.G.**, **H.B.**, **O.V.**, and **M.M.** wrote the manuscript with the contribution of all authors. All authors have approved the final version of the manuscript.

