## Supplementary Information for "Molecular trick to reverse S_N_2 mechanism in hydrolytic enzyme"

**Table S1:** Comparison of kinetic constants obtained for LinB haloalkane dehalogenase variants. Kinetic constants were obtained after mixing the chloride anion (Cl^-^) product or the 1-chlorohexane substrate ligands with LinB-wt, LinB-H272F, or LinB-H272N at either pH 7.5 or 10.5. The values are represented as best fit values ± standard error values based on nonlinear curve fitting. The details of kinetic data analysis are provided in **Supplementary Note 1**. n. a. = not applicable

|  | *K*_d_ (Cl^-^) [mM] | *K*_s_ [µM] | *k*_+2_ [s^-1^] | *k*_-2_ [s^-1^] | *K*_+2_ (*k*_+2_/ *k*_-2_) [–] |
| --- | --- | --- | --- | --- | --- |
| LinB-wt  (pH 7.5) | 620 ± 160 | 90 ± 10 | 84 ± 3 | 33 ± 1 | 2.55 ± 0.12 |
| LinB-wt  (pH 10.5) | 470 ± 80 | 180 ± 20 | 63 ± 1 | 33 ± 1 | 1.91 ± 0.07 |
| LinB-H272F  (pH 7.5) | 340 ± 80 | 540 ± 80 | 0.025 ± 0.002 | 0.074 ± 0.001 | 0.34 ± 0.03 |
| LinB-H272F  (pH 10.5) | 250 ± 60 | 250 ± 40 | 0.069 ± 0.037 | 0.137 ± 0.075 | 0.50 ± 0.39 |
| LinB-H272N  (pH 7.5) | 370 ± 140 | 320 ± 70 | 6.8 ± 0.4 | 0.007 ± 0.006 | 1010 ± 520 |
| LinB-H272N  (pH 10.5) | 240 ± 70 | 690 ± 110 | < 0.01 | < 0.01 | n. a. |

**Table S2:** Crystallographic data collection and refinement statistics.

| **Data collection*** | **LinB-H272F** |
| --- | --- |
| Wavelength (Å) | 1 |
| Space group | *P*2_1_2_1_2_1_ |
| Cell dimensions |  |
| a, b, c (Å) | 51.496, 65.448, 90.171 |
| α, β, γ (°) | 90, 90, 90 |
| Resolution (Å) | 44.17 – 1.55 (1.61 – 1.55) |
| Total reflections | 586,446 (56,284) |
| Unique reflections | 44,614 (4,309) |
| Rmerge | 18.0 (160.1) |
| I / σI | 11.71 (1.47) |
| Completeness (%) | 99.4 (97.6) |
| Multiplicity | 13.1 (13.1) |
| CC(1/2) | 99.8 (70.1) |
| Wilson B-factor | 13.09 |
| **Refinement** |  |
| Resolution (Å) | 44.172 – 1.551 (1.607 – 1.551) |
| No. reflections | 44,614 (4,309) |
| Rwork (%) / Rfree (%) | 17.41 / 20.13 |
| Number of atoms |  |
| Protein | 2,390 |
| Ligand | 102 |
| Water | 246 |
| B-factors |  |
| Protein | 14.6 |
| Ligand | 29.5 |
| Water | 26.7 |
| R.m.s deviations |  |
| Bond lengths (Å) | 0.007 |
| Bond angles (º) | 0.96 |
| Ramachandran favoured (%) | 96.23 |
| Ramachandran allowed (%) | 3.77 |
| Ramachandran outliers (%) | 0 |
| PDB ID code | 7NFZ |
| *Values in parentheses are for the highest-resolution shell. | |

**Table S3:** Residues used for the setup of the ASMD simulations.

| **Protein** | **Tunnel** | **Steering residue** |
| --- | --- | --- |
| LinB-H272F | p1 | Tyr 82 |
|  | p2 | Pro 203 |
|  | p5 | Ser 206 |
| LinB-H272N | p1 | Leu 89 |
|  | p2 | Met 21 |
|  | p5 | Arg 201 |
| LinB-wt | p1 | Tyr 82 |
|  | p2 | Arg85 |
|  | p5 | Glu 161 |

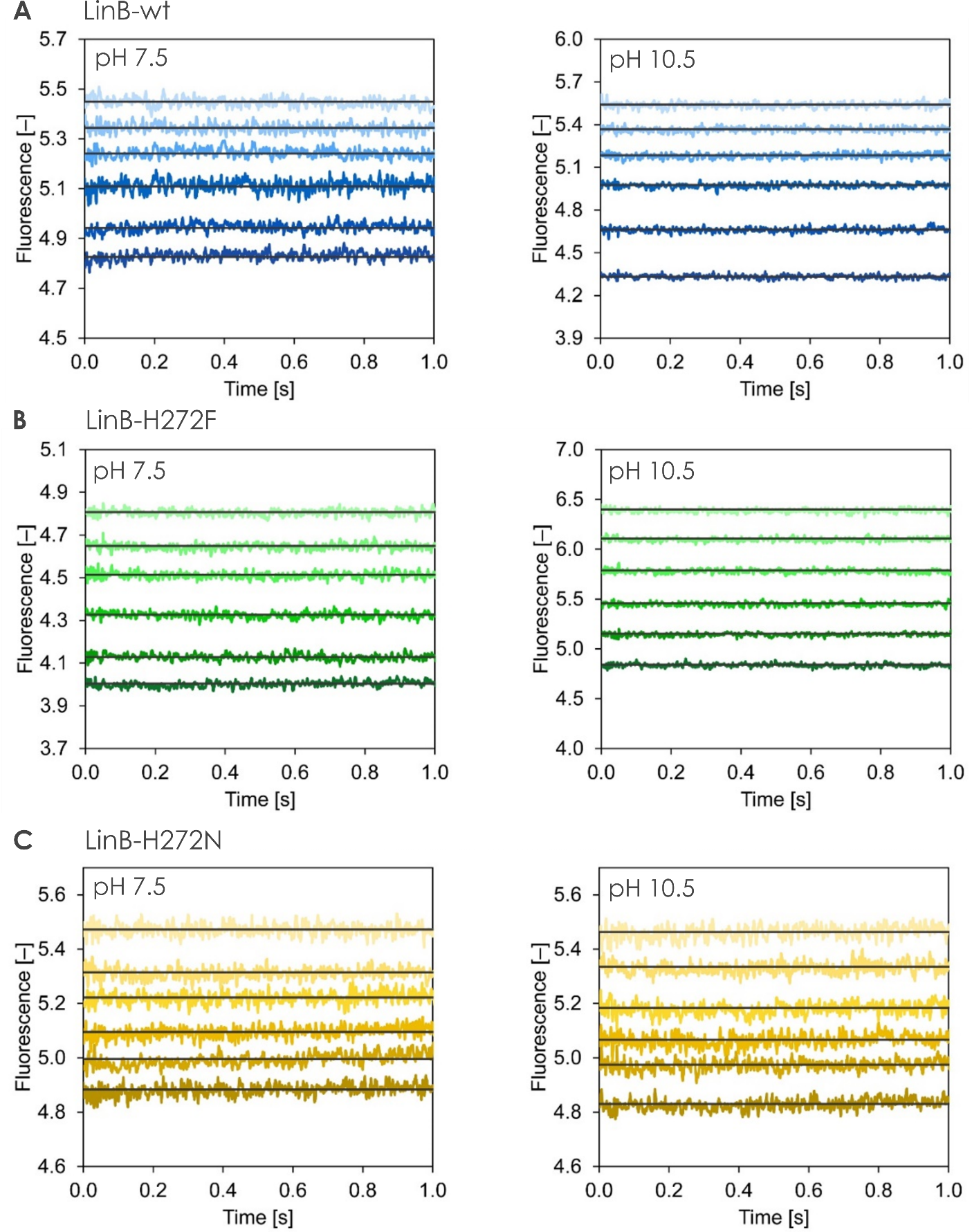

**Figure S1:** **Kinetic traces of Cl^-^ binding to LinB variants:** LinB-wt (A), LinB-H272F (B), and LinB-H272N (C), at pH 7.5 (left) and pH 10.5 (right). Solid lines represent the best fit. The concentration range of Cl^-^ during the measurements was between 0 and 2000 mM.

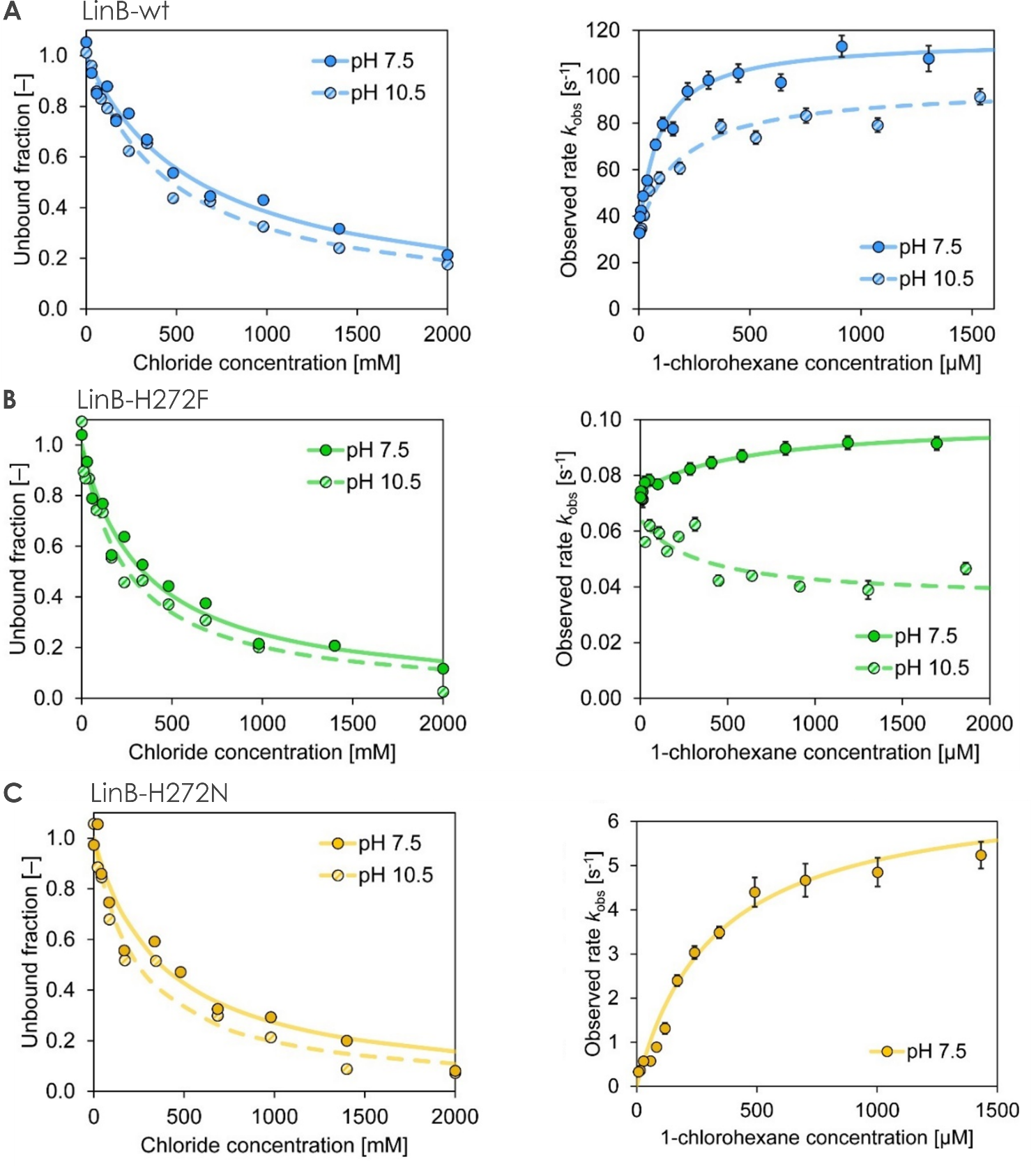

**Figure S2:** **Transient kinetic analysis of Cl^-^ (left) and 1-chlorohexane (right) for all three LinB variants:** LinB-wt (A), LinB-H272F (B), and LinB-H272N (C), at pH 7.5 (solid points) or pH 10.5 (hatched points). The analytical fitting of hyperbolic concentration dependencies was performed for equilibrium fluorescence levels upon Cl^-^ binding and for observed exponential rates, *k*_obs_, of fluorescence decrease upon 1-chlorohexane binding and processing. Lines represent the best fit.

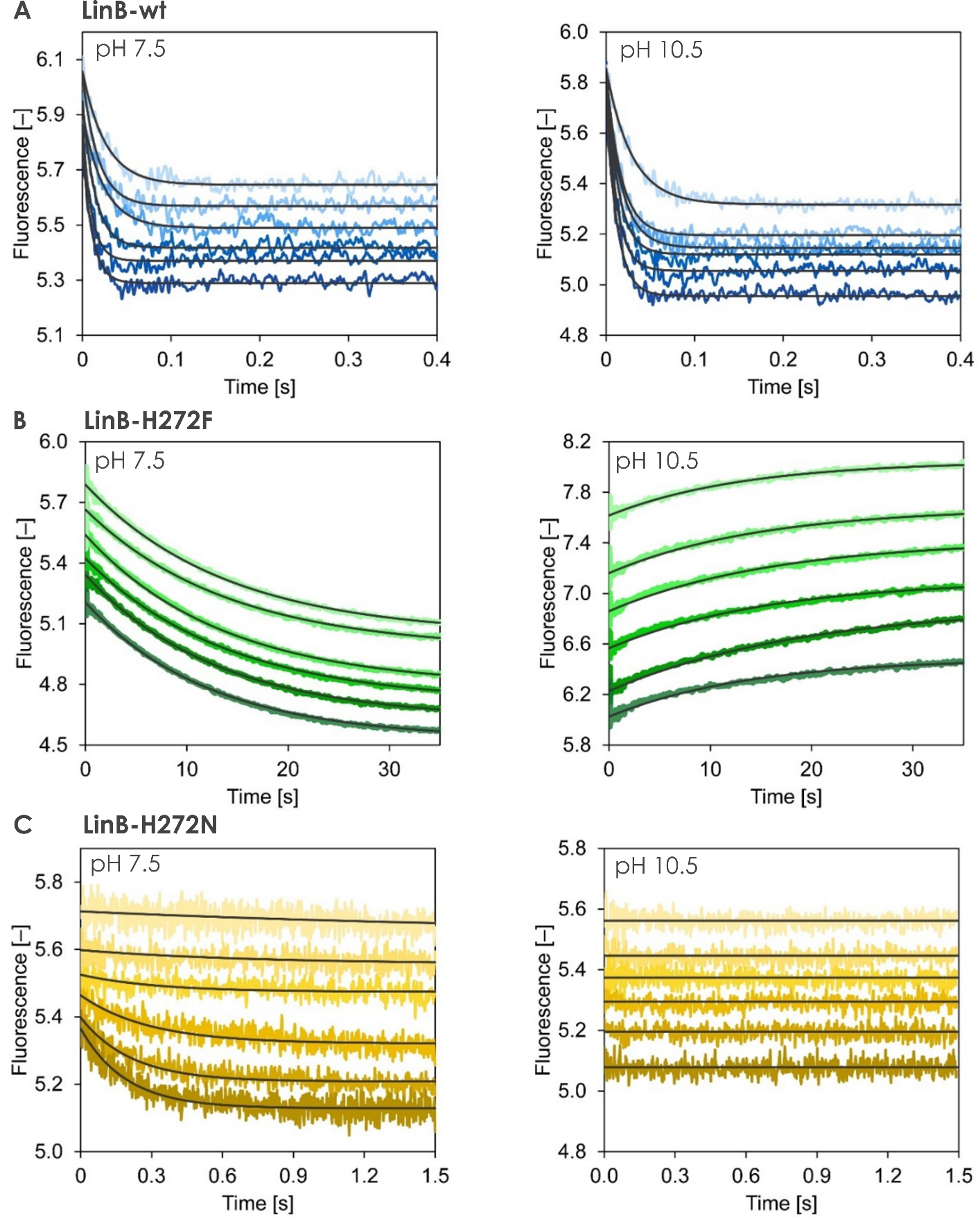

**Figure S3:** **Kinetic traces after mixing 1-chlorohexane (substrate) with LinB variants:** LinB-wt (A), LinB-H272F (B), and LinB-H272N (C), at pH 7.5 (left panels) and pH 10.5 (right panels). Solid lines represent the best fit. The concentration range of 1-chlorohexane during the measurements was between 0 and 1860 μM.

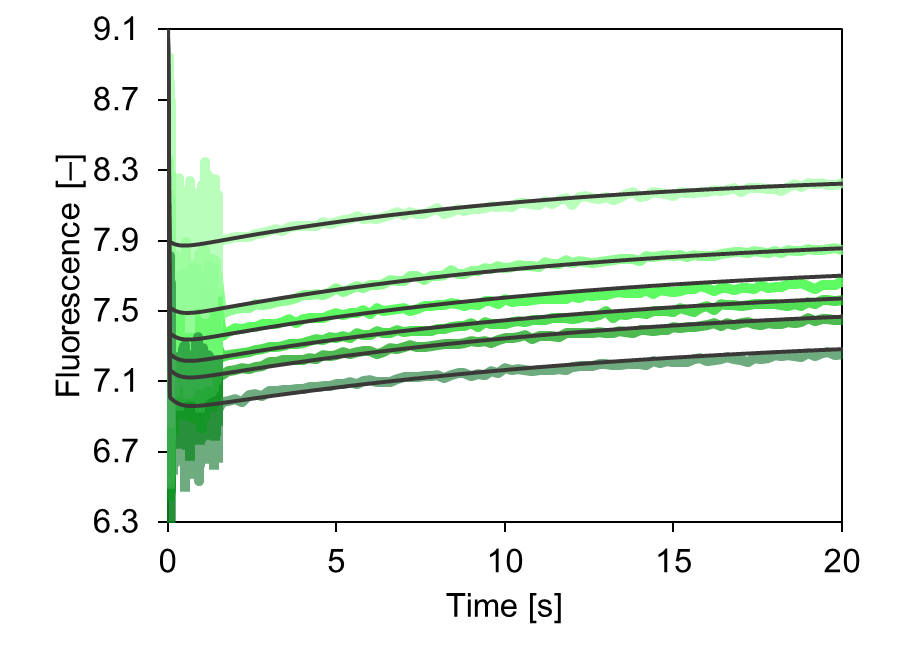

**Figure S4:** Numerical fitting of the LinB-H272F kinetic data upon mixing with 1-chlorohexane at pH 10.5. Due to the complex behaviour of the mutant at higher pH, including the conformational selection binding mechanism and chloride release after the alkyl-enzyme intermediate formation, numerical data analysis was required. The final fit (solid lines) could adequately account for all the untypical experimental observations, supporting the proposed extended model.

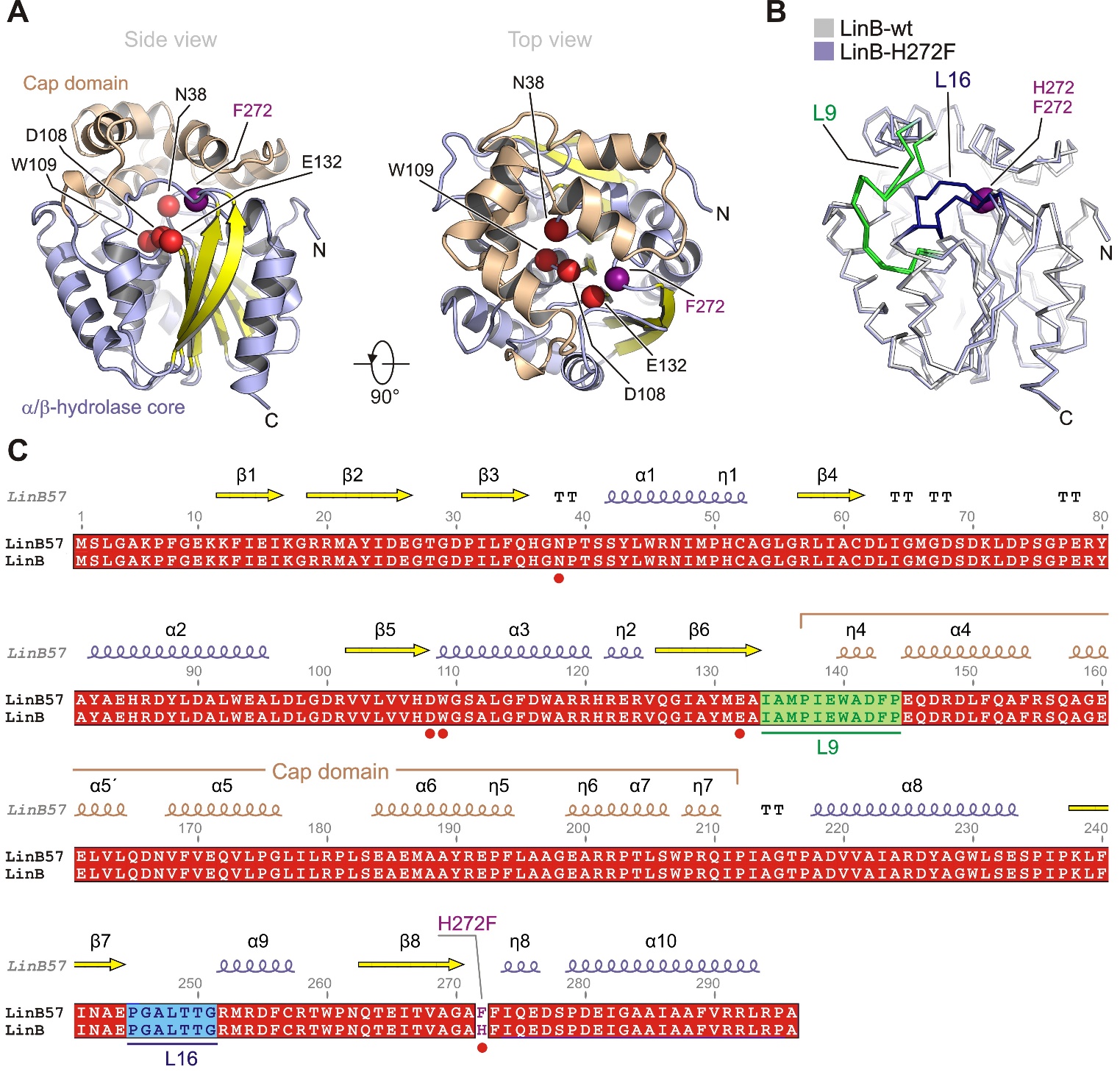

**Figure S5:** Overall structure of LinB-H272F. (A). Cartoon representation of LinB-H272F. Catalytic pentad residues are shown as red spheres, with the exception of introduced F272 that is colored violet. The central 8-stranded sheet is colored yellow, the helical cap domain is colored wheat, the remaining parts are colored light-blue. (B) Ribbon representation of structural comparison between LinB-wt (grey) and LinB-H272F (light-blue). The L9 loop is colored green, and the L16 loop is colored dark-blue. Note that major structural deviations are made by both L9 and L16 loops. (C) Sequence alignment between LinB-H272F and LinB-wt. Topology of secondary structure elements is denoted above the alignment.

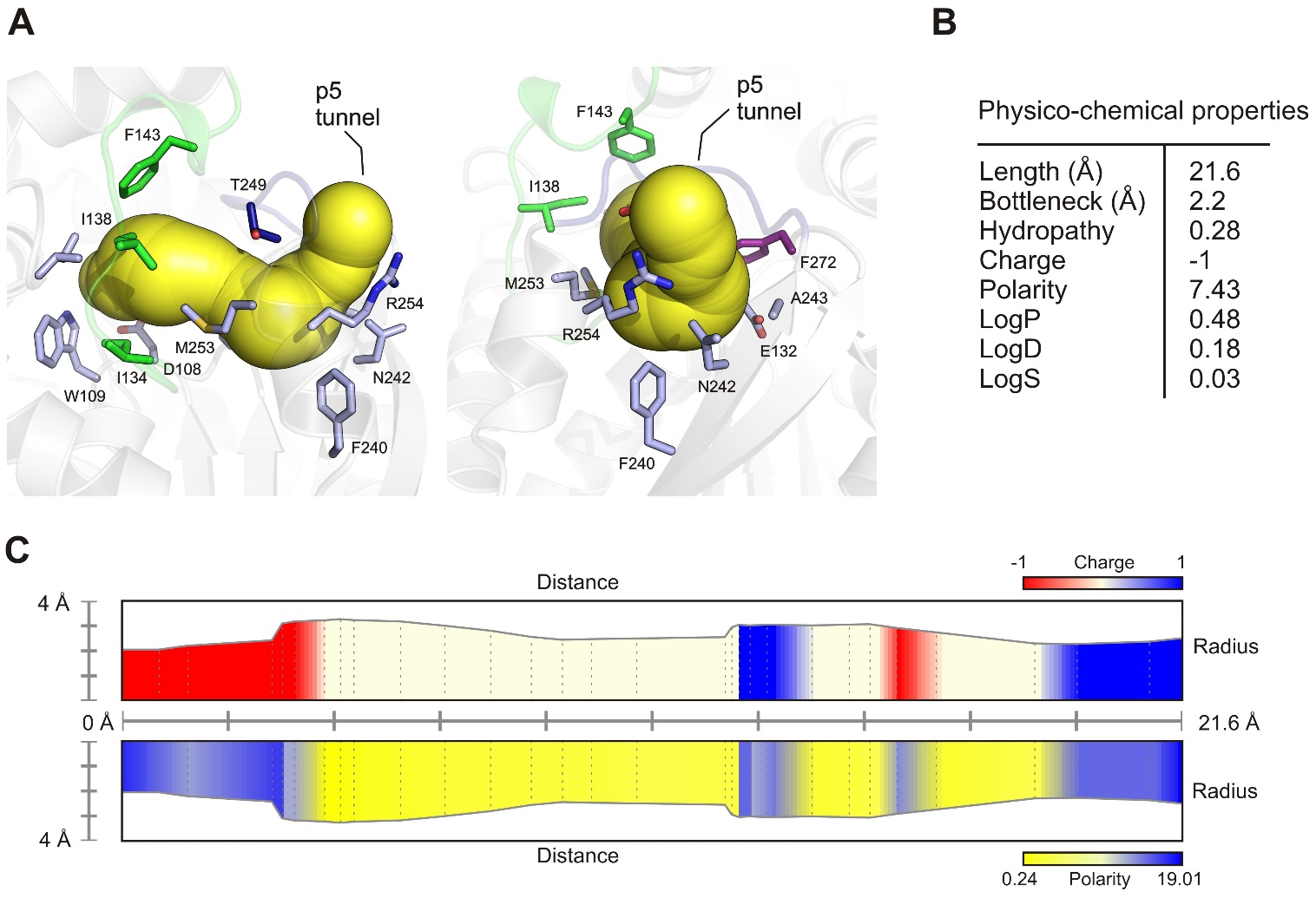

**Figure S6:** Analysis of the p5 tunnel in LinB-H272F. (A) Identification of vizualization of p5 tunnel (yellow) by MOLE software tool.^1^ Amino acids surrounding the tunnel are shown as sticks. (B) Table with physico-chemical properties of the p5 tunnel. (C) The charge and hydrophobicity properties of the p5 tunnel.

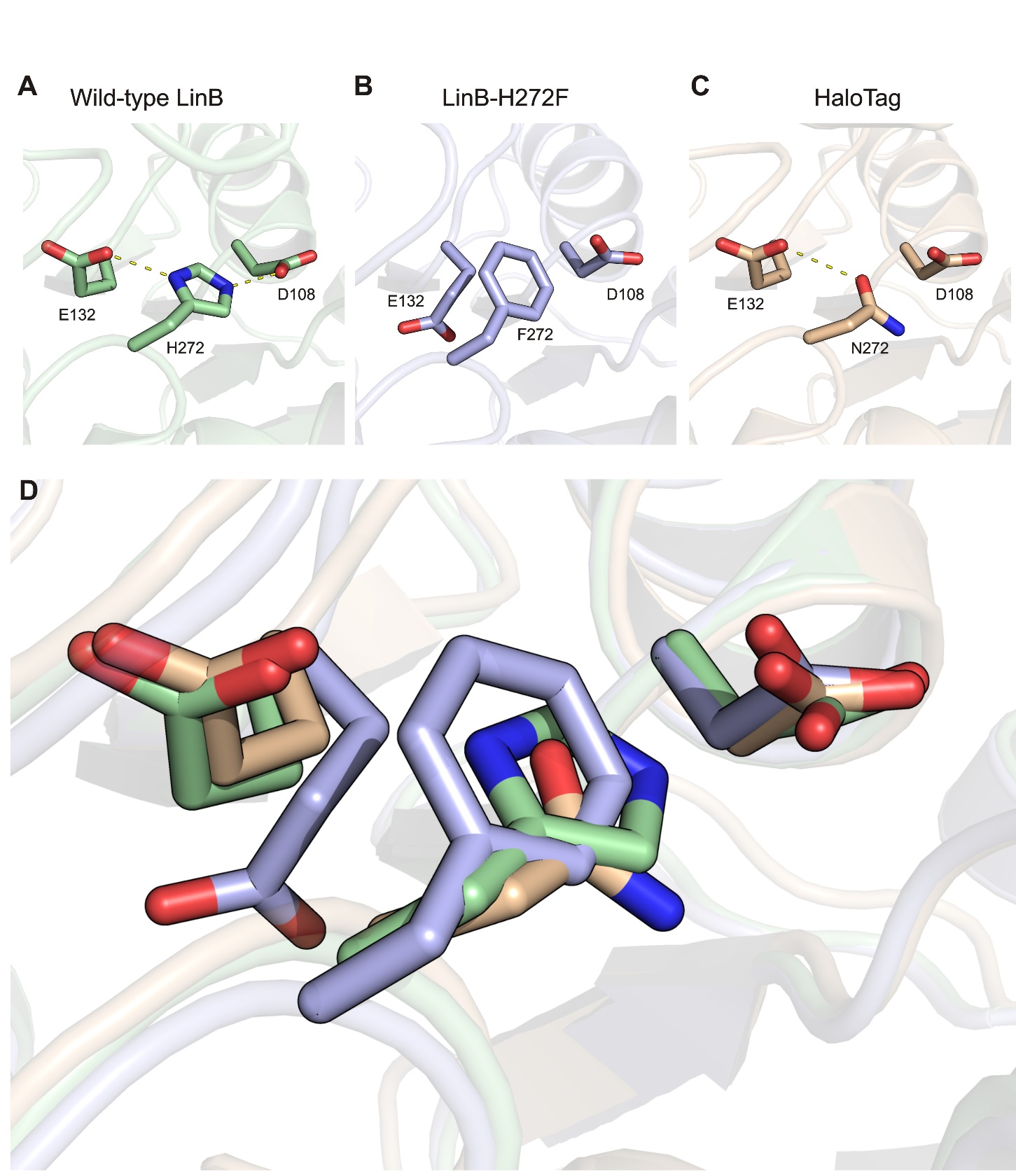

**Figure S7:** The positioning of catalytic triad residues (proton-relay system) in LinB-wt (A), LinB-H272F (B), and HaloTag (C). Note that a catalytic acid (E132) is flipped-out from its canonical position in LinB-H272F. (D) Superposition of all structures. LinB-wt (lightgreen), LinB-H272F (lightblue), and HaloTag (wheat).

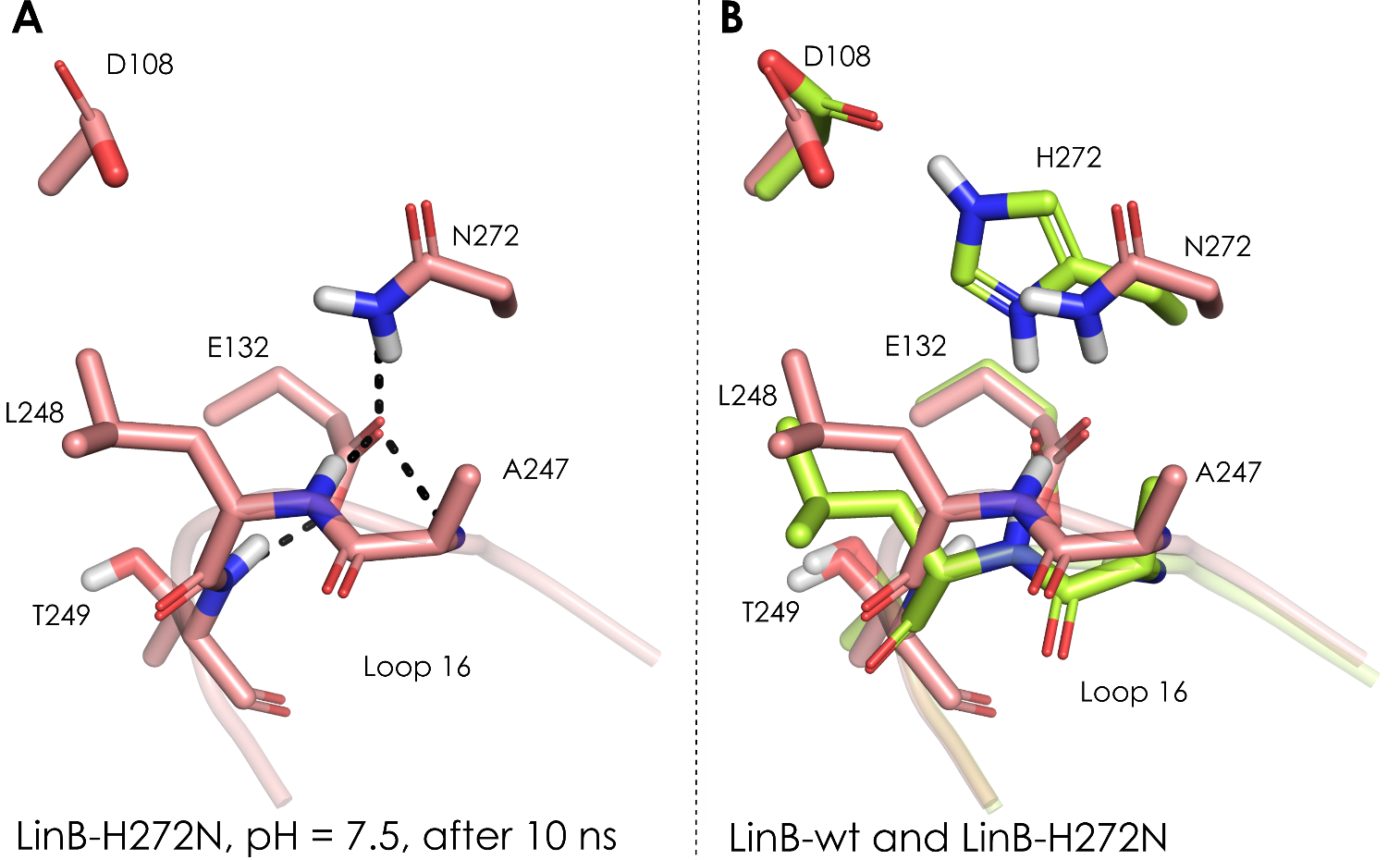

**Figure S8:** LinB-H272N after 10 ns of MD simulations. A: D108 conformation changes compared to the initial Rosetta model (see **Figure 5D** of the main article), resulting in the loss of the H-bond between D108 and N272. B: Superimposition of LinB-wt and LinB-H272N.

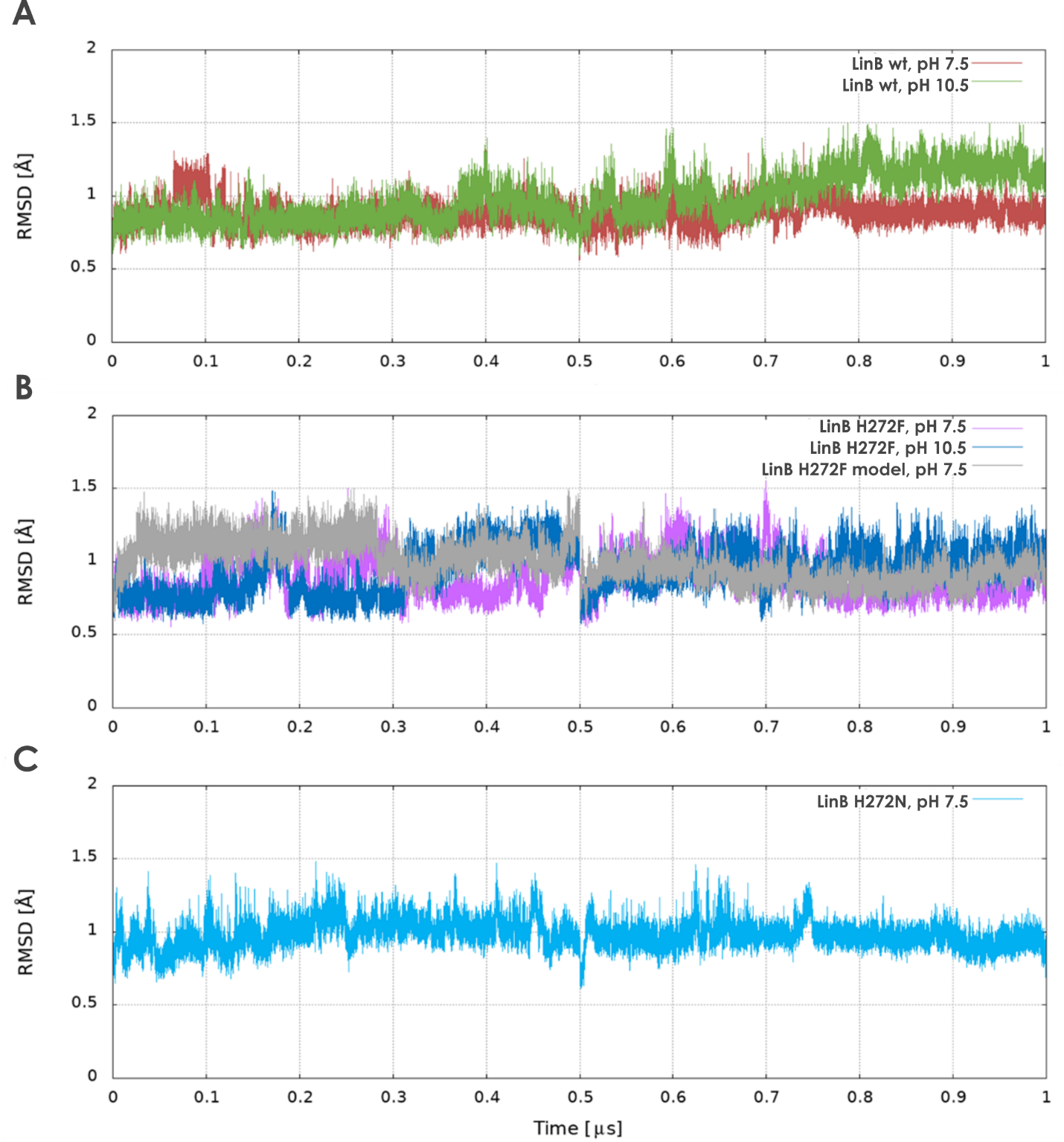

**Figure S9:** Variation of the root-mean-square deviation (RMSD) of the backbone atoms during the MD simulations with respect to the initial structures, for the different systems at different pH values. A: LinB-wt, B: LinB-H272F (started from the crystal structure and the *in silico* model), C: LinB-H272N (pH 7.5). Two independent MD simulations were performed for every protein, each one consisting of 500 ns. Here, they are presented sequentially. The plateaus observed at the final portion of every MD, together with the low RMSD values, below 1.5 Å, confirm their stability.

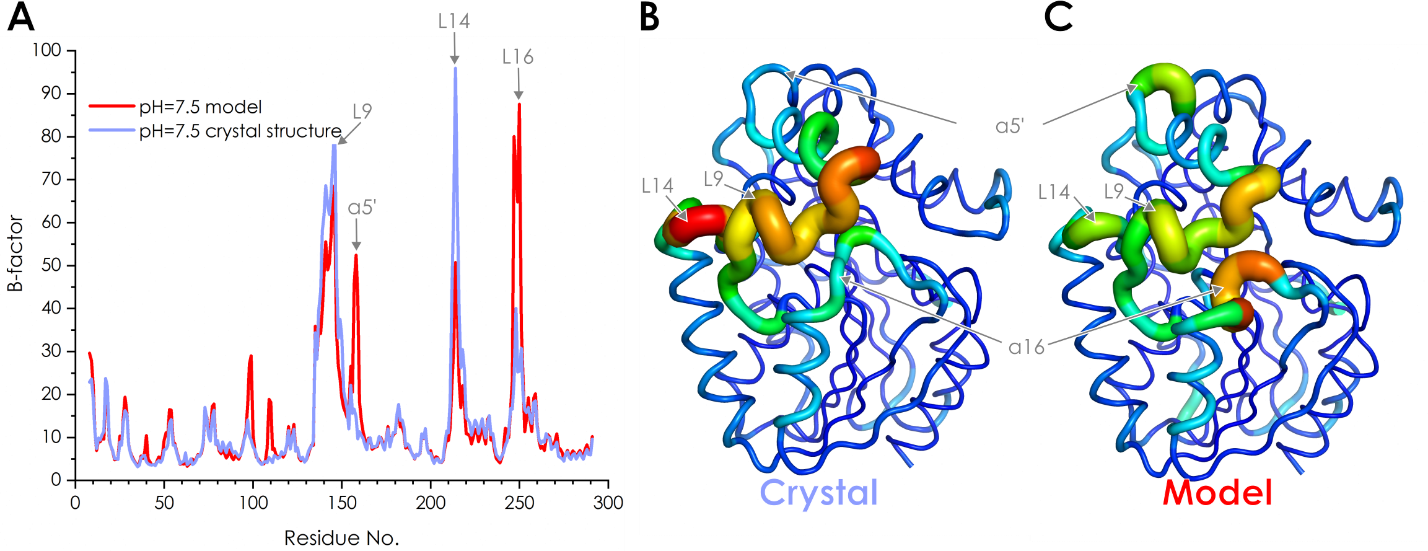

**Figure S10:** B-factor values during MD simulations of LinB-H272F at pH 7.5. A: Graph representation of B-factor values of crystal structure (light blue) and model (red). B and C: Putty tubes representation of crystal and model structure, respectively. A hotter (redder) and thicker putty tube indicates higher B-factor (higher flexibility); the regions with more pronounced flexibility changes are indicated.

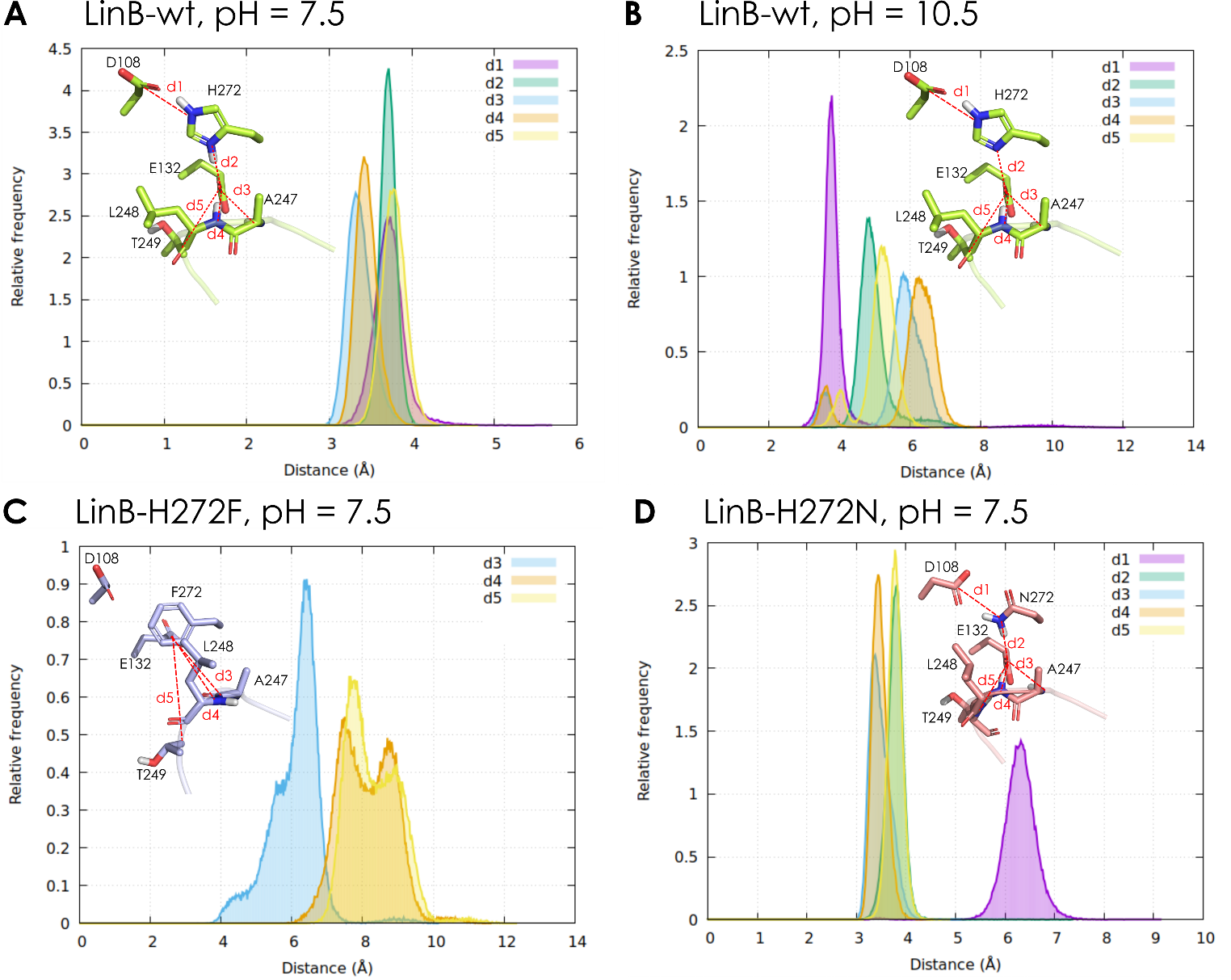

**Figure S11:** Distribution of interatomic distances during the MD simulations, representing the H-bond network shown in **Figure 5** of the main article. LinB-wt at pH 7.5 (A), LinB-wt at pH 10.5 (B), LinB-H272F at pH 7.5 (C), and LinB-H272N at pH 7.5 (D). The distance values of 3-4 Å correspond to H-bonds. The insets in each panel depict the nature of distances: (i) d1 (D108-Cγ–H272-Nε for LinB-wt or D108-Cγ–N272-Nδ for LinB-H272N), (ii) d2 (E132-Cδ–H272-Nδ or E132-Cδ–N272-Nδ), (iii) d3 (E132-Cδ–A247-N), (iv) d4 (E132-Cδ–L248-N), and (v) d5 (E132-Cδ–T249-N).

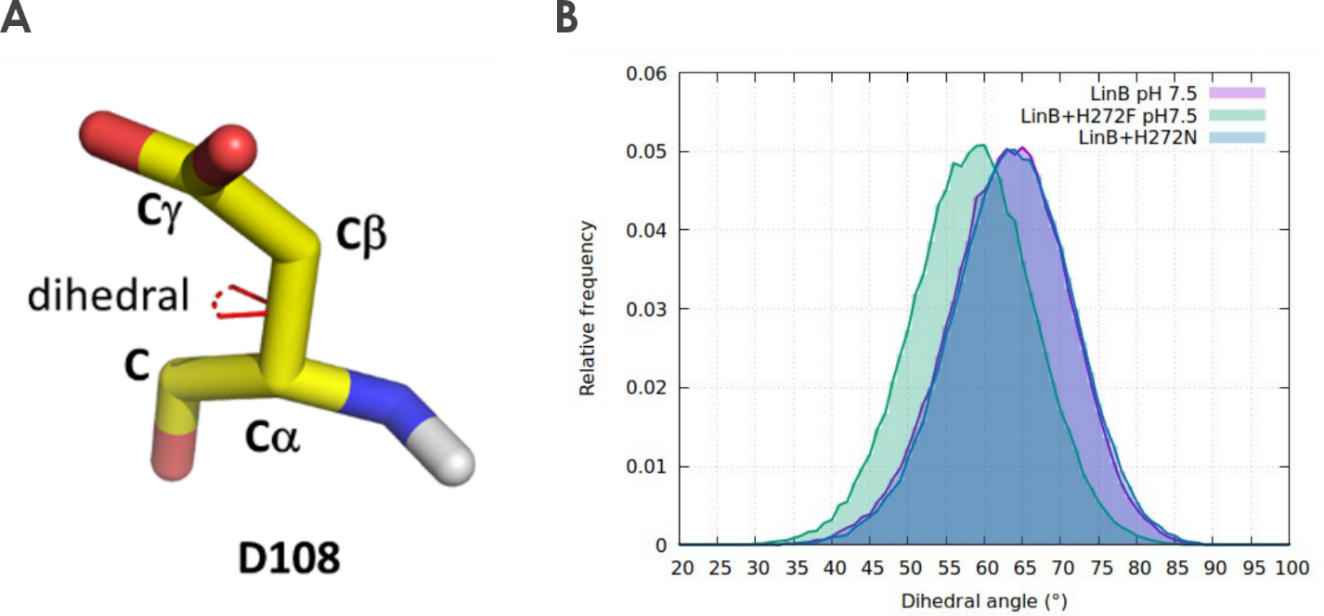

**Figure S12:** Dihedral angle of the catalytic D108. A: The dihedral angle of D108 is defined as an angle between two planes. First plane consists of atoms C-Cα-Cβ and the second plane consists of atoms Cα-Cβ-Cγ. The dihedral angle also corresponds to the rotation around the two atoms, Cα and Cβ, as depicted in the figure. B: Distribution of the D108 dihedral angle in the MD simulations at pH 7.5 for LinB-wt (purple), LinB-H272F (green), and LinB-H272N (blue).

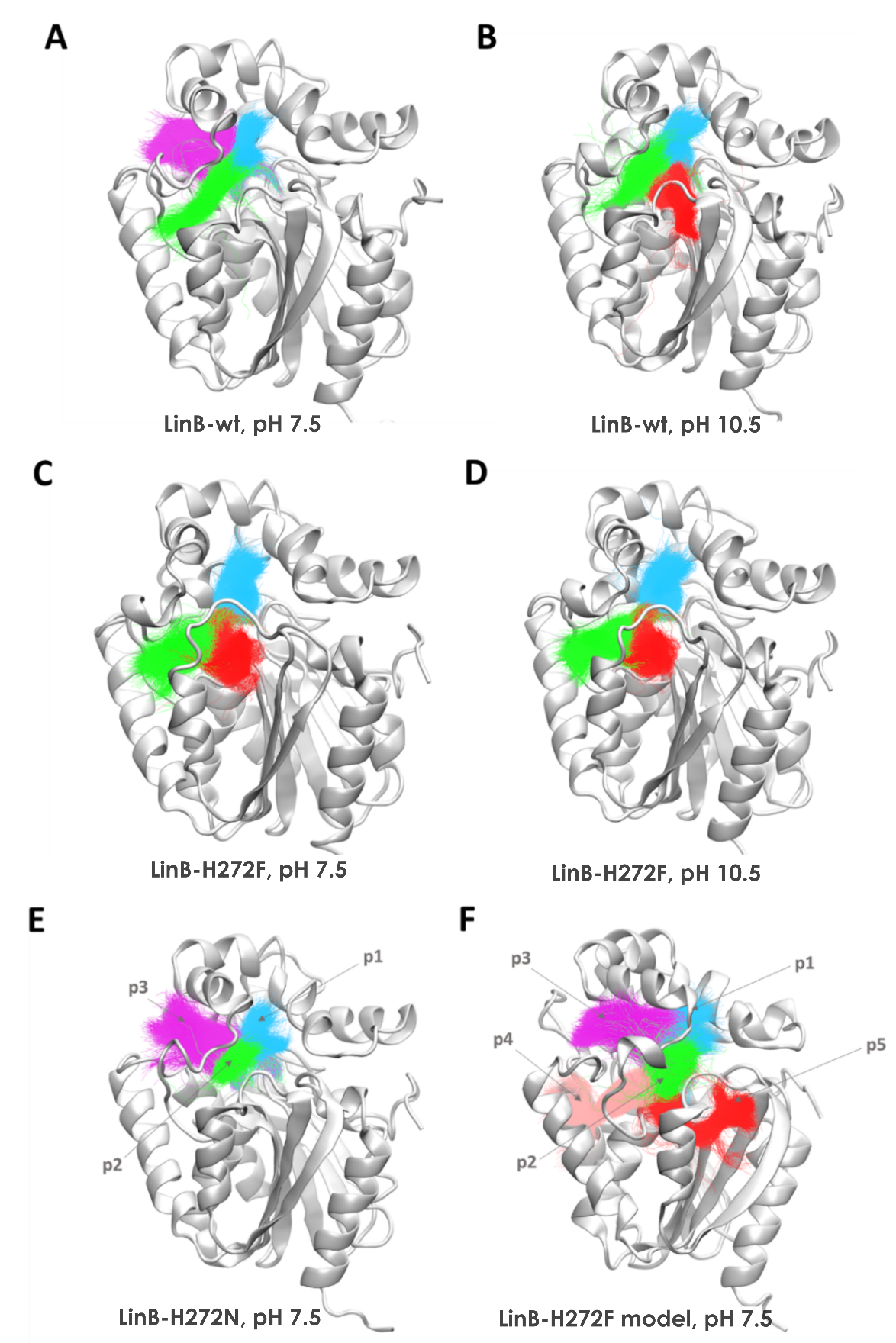

**Figure S13:** Access tunnels calculated in the combined MD simulations for LinB-wt (A and B), and LinB-H272F (C and D) at neutral (left) and alkaline (right) pH, LinB-H272N (E), and the *in silico* model of LinB-H272F (F) at pH 7.5. The tunnels are represented by the superimposed coloured lines, collected from the simulation snapshots, and clustered according to their topologies. p1 (cyan), p2 (slot tunnel, green), p3 (magenta), p4 (salmon) p5 (red).

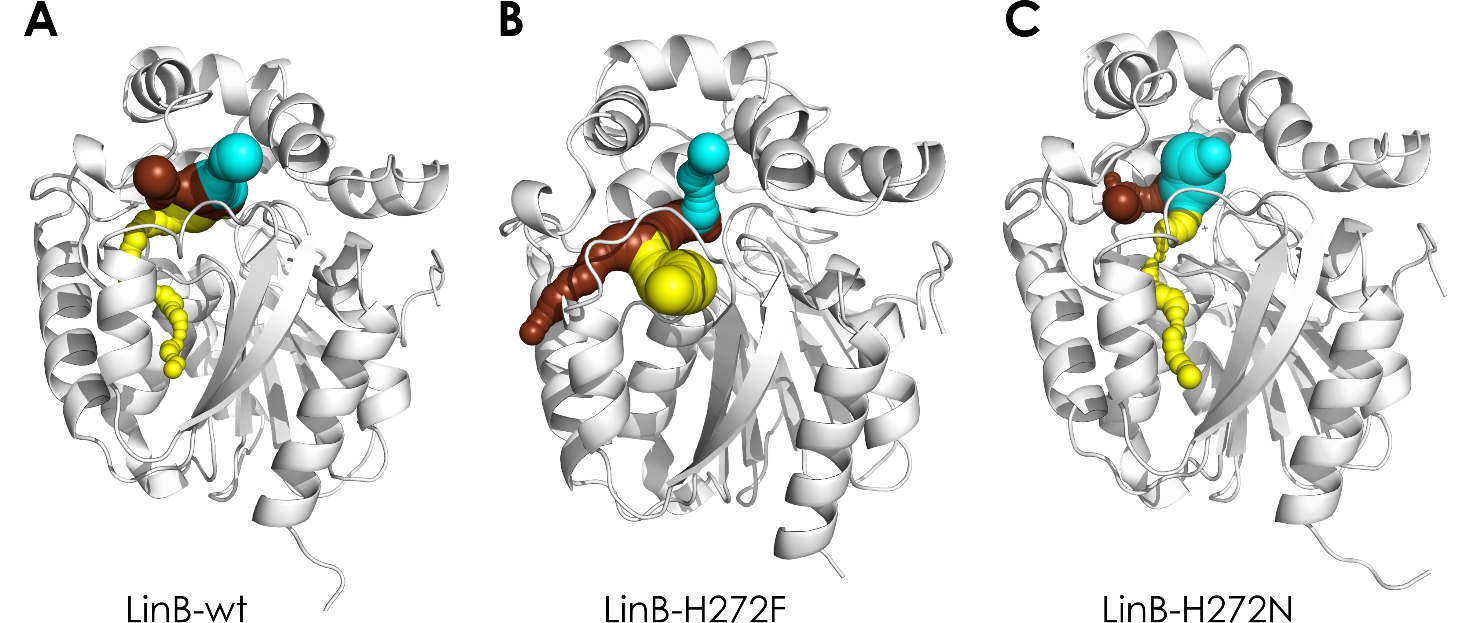

**Figure S14:** Structures of LinB variants with calculated tunnels using the probe with radius of 0.7 Å. LinB-wt (A), LinB-H272F (B), and LinB-H272N (C). p1 tunnel (cyan), p2 tunnel (brown), and p5 tunnel (yellow).

**Supplementary Note 1:**

**Kinetic analysis of 1-chlorohexane processing by LinB variants**

Apart from the typical single-exponential decrease of fluorescence traces with the increasing hyperbolic concentration dependence of observed exponential rates *k*_obs_ (described in the Materials and Methods section), more complex and untypical kinetic data were collected in the case of histidine mutants LinB-H272F and LinB-H272N at the increased value of pH 10.5.

When LinB-H272F was mixed with 1-chlorohexane at increased pH of 10.5, (i) the fluorescence signal was unexpectedly increasing with time after a quick signal drop in the instrument dead-time, and furthermore, (ii) the concentration dependence of observed rates yielded a decreasing trend. The increase of the fluorescence signal and regeneration to the original level indicated that the chloride anion was released from the enzyme active site after the alkyl-enzyme intermediate was formed. This is unique in comparison with LinB-wt, other common haloalkane dehalogenases, and even LinB-H272F at physiological pH where the halide anion does not leave until the alkyl-enzyme intermediate is hydrolyzed and both the alcohol as well as the halide products are released.

The hyperbolic decrease of the observed rate with increasing substrate concentration pointed out the conformational selection mechanism of the substrate binding by LinB-H272F at pH 10.5. None of these two events has ever been observed previously in haloalkane dehalogenases and it is presumed that the untypical behavior might be due to the change of protonation states at alkaline pH causing a loss of important residue contacts and a newly formed tunnel which was detected in crystal structures and MD simulations of this variant. This extended model of substrate processing was tested by numerical simulation data fitting and a good fit, accounting for the unusual observations, could be obtained (**Figure S14**). This result supported the postulated hypothesis, but further experiments would be needed to fully confirm the suggested mechanism.

Finally, the kinetics of 1-chlorohexane processing by LinB-H272N was also affected by the increased pH environment. Fluorescence traces exhibited no significant decrease within the experimental time window of 100 seconds, suggesting highly impaired kinetics with the values of exponential observed rates lower than 0.01 s^-1^. This value set the upper limit for both the forward and reverse rate constants of the alkyl-enzyme intermediate formation (*k*_+2_ and *k*_-2_) without a possibility to determine exact values which were too slow to be derived from the obtained kinetic traces. The value of the substrate binding dissociation constant *K*_s_ was derived based on the concentration dependence analysis of the drop of the initial fluorescence level caused by the bound substrate, showing a typical hyperbolic dependence.
